# Molecular recognition and dynamics of linear poly-ubiquitins: integrating coarse-grain simulations and experiments

**DOI:** 10.1101/2020.04.14.041327

**Authors:** Alexander Jussupow, Ana C. Messias, Ralf Stehle, Arie Geerlof, Sara M. Ø. Solbak, Anders Bach, Michael Sattler, Carlo Camilloni

## Abstract

Poly-ubiquitin chains are flexible multidomain proteins, whose conformational dynamics enable their molecular recognition by a large number of partners in multiple biological pathways. By using alternative linkage, it is possible to obtain poly-ubiquitin molecules with different dynamical properties. This flexibility is further increased by the possibility to tune the length of poly-ubiquitin chains. Characterizing the dynamics of poly-ubiquitins as a function of their length is thus relevant to understand their biology. Structural characterization of poly-ubiquitin conformational dynamics is challenging both experimentally and computationally due to increasing system size and conformational variability. Here, by developing highly efficient and accurate small-angle X-ray scattering driven Martini coarse-grain simulations, we characterize the dynamics of linear M1-linked di-, tri- and tetra-ubiquitin chains. Our data show that the behavior of the di-ubiquitin subunits is independent of the presence of additional ubiquitin modules. We propose that the conformational space sampled by linear poly-ubiquitins, in general, may follow a simple self-avoiding polymer model. These results, combined with experimental data from small angle X-ray scattering, biophysical techniques and additional simulations show that binding of NEMO, a central regulator in the NF-κB pathway, to linear poly-ubiquitin obeys a 2:1 (NEMO:poly-ubiquitin) stoichiometry in solution, even in the context of four ubiquitin units. Eventually, we show how the conformational properties of long poly-ubiquitins may modulate the binding with their partners in a length-dependent manner.

**Significance:** Protein conformational dynamics plays an essential role in molecular recognition mechanisms. The characterization of conformational dynamics is hampered by the conformational averaging of observable in experimental structural biology techniques and by the limitations in the accuracy of computational methods. By developing an efficient and accurate approach to combine small-angle X-ray scattering solution experiments and coarse-grain Martini simulations, we show that the conformational dynamics of linear poly-ubiquitins can be efficiently determined and to rationalize the role of poly-ubiquitin dynamic in the molecular recognition of the UBAN domain upon binding to the signaling regulator NEMO. The analysis of the conformational ensembles allows us to propose a general model of the dynamics of linear poly-ubiquitin chains where they can be described as a self-avoiding polymer with a characteristic length associated with their specific linkage.

## Introduction

Ubiquitination is a reversible post-transcriptional modification system that regulates key physiological processes, such as protein degradation, cell cycle, apoptosis, DNA repair, and signal transduction^1-3^. Once a protein substrate is mono-ubiquitinated (e.g. a lysine of the substrate is conjugated through an iso-peptide bond to the C-terminus of a ubiquitin monomer), an additional ubiquitin may be conjugated to either one of the seven lysine residues of the first ubiquitin (K6, K11, K27, K29, K33, K48, and K63)^4^ or its amino-terminal methionine residue (M1)^5-7^. This process can lead to the assembly of poly-ubiquitin chains of various lengths and topologies. The resulting polymeric chains are then associated with different cellular mechanisms^8^. Since all these polymers are made of the same single unit, the highly conserved 76-residues long ubiquitin domain, the ubiquitin code is an example of a conformation-based alphabet, where both the polymerization site^8-9^ as well as the chain length^10^ regulate the recognition by different partners, and thereby determine the cellular fate of the protein. The role of poly-ubiquitin length and dynamics in molecular recognition processes is poorly understood^8, 10-11^. An overall assessment of the typical length of different poly-ubiquitin chains in physiological conditions is missing, and only sporadic indications are available. For example, in the case of K48-linked poly-ubiquitin, a length of four is generally considered optimal for molecular recognition of the 26S proteasome^12^, while the nuclear protein localization protein 4 (Npl4) is selective for K48-linked chains longer than six^13^. Interestingly, it was reported that K48-linked tetra-ubiquitin (Ub_4_) slows down further ubiquitination^14-16^, while this is not the case for K63-linked Ub_4_^16^.

Linear M1-linked poly-ubiquitin chains (**Figure 1**), whose assembly is catalyzed by LUBAC^5^, have been shown to play a role in inflammation, immune responses, and oncogenesis^17-19^. Their most studied function is the involvement in the activation of the canonical NF-*κ*B pathway^6-7, 17, 20-23^. In this pathway, the IKK complex (or I*κ*B kinase, formed by IKK*α*, IKK*β*, and NEMO, also known as IKK*γ*, the NF-*κ*B essential modulator) is activated by LUBAC upon activation by various stimuli^22^. LUBAC preferentially recognizes and conjugates linear ubiquitin chains on NEMO. NEMO also possesses a specific linear di-ubiquitin-binding region referred to as the “ubiquitin binding in ABIN and NEMO” (UBAN) motif^24^, which forms a helical coiled-coil dimer in solution^23^. Recognition of a linear poly-ubiquitin conjugated to NEMO by the UBAN domain of another NEMO may trigger the clustering of the IKK complex as well as conformational changes that subsequently activate IKK^25-26^. Once active IKK can phosphorylate and inactivate the I*κ*Bs (inhibitor of the NF-*κ*B proteins) leading to the release of NF-*κ*B^27^. Indeed, it was recently shown that it is possible to inhibit NF-*κ*B activation upon UBAN-dependent TNFα and TCR/CD28 stimulation by small-molecules that inhibit the binding of linear poly-ubiquitins to the NEMO_UBAN_ domain^23^. While the NEMO_UBAN_ domain can bind linear di-ubiquitin, it has been observed that full-length NEMO can only bind Ub_4_ or longer suggesting a length-dependent activation mechanism^21^. Furthermore, another study suggested that the binding of NEMO to chains of 10 linear ubiquitins or longer induces a different conformation of NEMO compared to the binding of shorter chains^20^.

**Figure 1.**
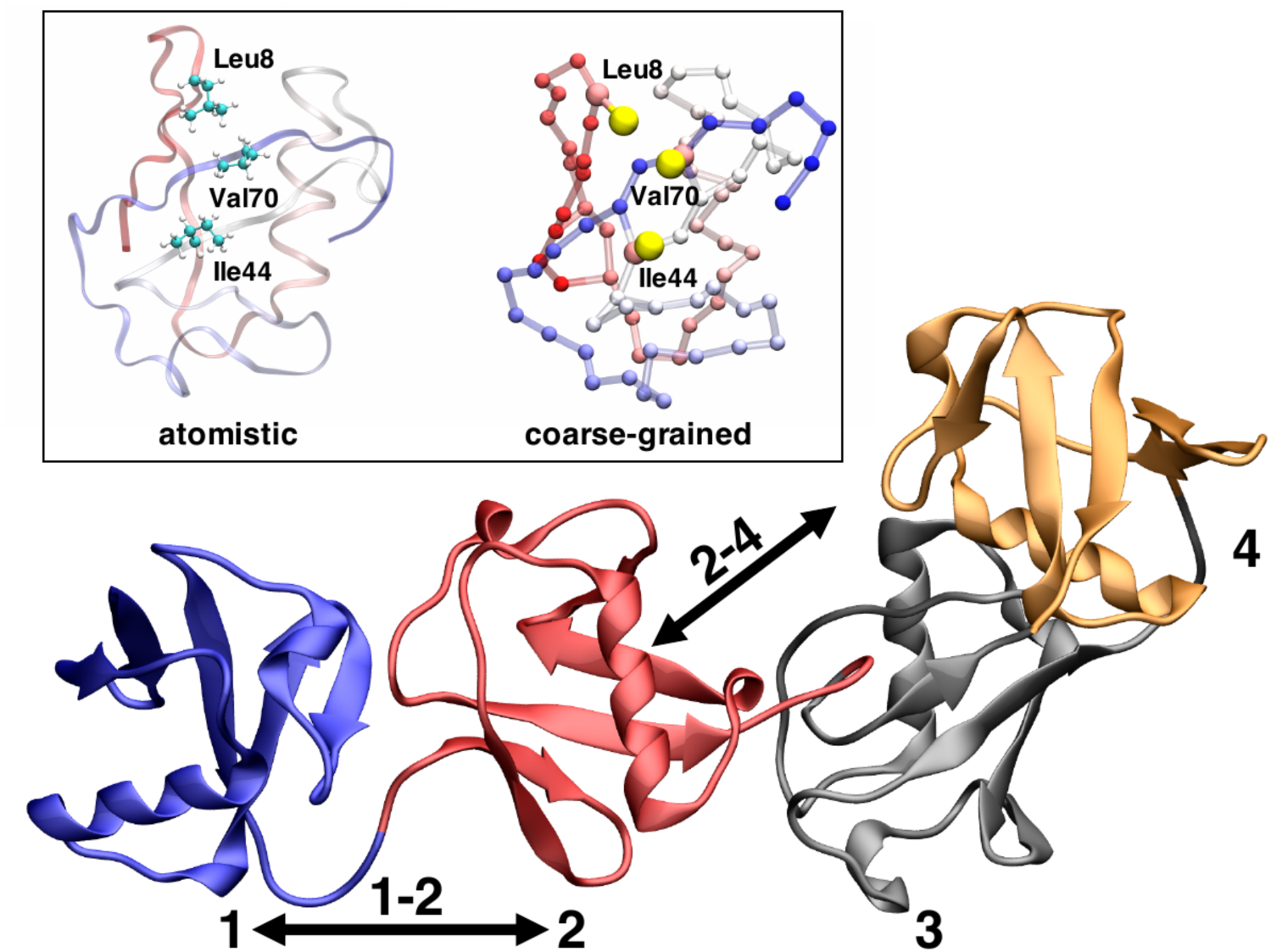
Schematic illustrations linear poly-ubiquitins. Cartoon representation of linear tetra-ubiquitin, the ubiquitin domains are numbered from the N-terminal to the C-terminal from 1 to 4. In the inset is shown an atomistic and coarse grained (Martini) ubiquitin domain highlighting the hydrophobic patch (Ile44, Val70 Leu8).

Characterizing the conformational space of poly-ubiquitin chains as a function of length is critical to understand their physiological behavior. Such structural characterization is nonetheless very challenging. Poly-ubiquitins, from di-ubiquitin to longer chains, exhibit a very dynamic behavior^28^ that requires determining a statistical ensemble of all the relevant configurations populated in solution. The combination of molecular dynamics (MD) with experimental small-angle X-ray scattering (SAXS) data is very well suitable to study dynamic protein systems^29^ including poly-ubiquitin of varying chain size. SAXS does not provide high-resolution structural information. Conversely, MD simulations may be used to determine the statistical ensemble of configuration populated by a system in equilibrium condition, but a full modeling base on MD simulations is hampered by the size of the system^30-31^. This problem can, in principle, be alleviated by coarse-grain force-fields^32^, eventually combined with enhanced sampling techniques^33^, that can massively speed up MD simulations although potentially at the expense of the accuracy^32^.

Here, we show that by integrating SAXS and MD simulations based on the Martini coarse-grain force-field^34-35^ by means of Metainference^36^ we can efficiently generate an ensemble of structures representing the dynamics of linear poly-ubiquitins (**Figure 1**). The ensembles allow the description of the dynamics of such complex systems at the single residue level. Our results show how poly-ubiquitins can populate multiple conformational states, but unexpectedly indicate that linear poly-ubiquitin chains and potentially poly-ubiquitins, in general, can be described by a simple self-avoiding polymer model. Various biophysical experiments are used to characterize the stoichiometry, kinetics and thermodynamic properties of the binding of poly-ubiquitin to the NEMO_UBAN_ domain. Surprisingly, our data demonstrate that NEMO_UBAN_ binds to di-, tri- and tetra-ubiquitin (Ub_2_, Ub_3_, and Ub_4_) in all cases forming a 2:1 NEMO_UBAN_:Ub_N_ complex in solution. Notably, a conformational ensemble for the NEMO_UBAN_:Ub_2_ complex rationalizes the 2:1 binding. Combined with our proposed poly-ubiquitin polymer model, this suggests how longer poly-ubiquitin chains may modulate NEMO recognition as well as bind more than one NEMO dimer.

## Results and Discussion

### A simple Martini modification improves the simulation of linear di-ubiquitin

We first evaluated the ability of the Martini coarse-grain force field to describe the dynamics of a linear Ub_2_. A metadynamics^37^ simulation of Martini Ub_2_ resulted in an extremely compact ensemble of structures (**Figure 2a**) which does not reproduce the measured SAXS intensities (**Figure 2d** and **Fig. S1** in the Supporting Information). In **Figure 2a** we report a free energy landscape (in kJ/mol) as a function of the distance between the centers of the two ubiquitin domains and their relative orientation; the average distance between the two domains is very short, around 2.41 ± 0.02 nm, with a preferential orientation of the two ubiquitin’s domain (measured as the torsion angle between two axes defined using the first and second half of the sequence of each ubiquitin, cf. Methods). The average radius of gyration of 1.73 ± 0.01 nm, strongly underestimates the value of 2.23 ± 0.02 nm derived from SAXS (**Table S1**). The ensemble seems to be able only to capture compact Ub_2_ configurations also when compared to the available crystal structures (PDB 2W9N^38^ (open), 3AXC^39^ (compact) and 4ZQS^28^ (compact)). This result can indicate an imbalance between the protein-protein and the protein-solvent interaction in Martini, a result that is not unexpected^40-41^ given the extremely simple description employed for the solvent (more complex descriptions, like the Martini polarizable water^42^, are available at the expense of performances). Interestingly, recent developments in atomistic force fields demonstrated the need for tuning solute-solvent interactions^43-44^. Following recent approaches that have successfully improved atomistic and coarse-grained force-fields, we repeated the same simulation after increasing by 5% the Martini water-protein Lennard-Jones interaction. This simple adjustment was sufficient to obtain a more expanded ensemble of structures as shown by the free energy landscape (**Figure 2b**) without any additional computational cost (**Table S2**). Importantly, the new ensemble resulted in an improved, even if not yet quantitative, agreement with the SAXS data (**Figure 2d**, blue curve; **Fig. S1**). The average distance between the domains increased to 3.10 ± 0.02 nm, and the protein can explore a much wider conformational space that now includes open and closed structures. In terms of the radius of gyration, the ensemble average resulted in 2.05 ± 0.01 nm to be compared with the 2.23 ± 0.02 nm derived from SAXS. Of notice, an under-development version of the Martini force field (Martini 3, currently in beta phase) while showing promising behavior may still benefit from increased protein-water interaction (**Fig. S2**). Importantly, Martini simulations outperform full atomistic simulations for the same system. As reported in **Table S1** an explicit solvent atomistic Ub_2_ simulation is around 500 times slower than a Martini simulation. Nonetheless, our aim here is to obtain ensembles in quantitative agreement with the SAXS data without a large-scale force field re-parameterization effort. This can be achieved at least, in principle, by integrating experimental information directly in the simulation by Metainference^36^.

**Figure 2.**
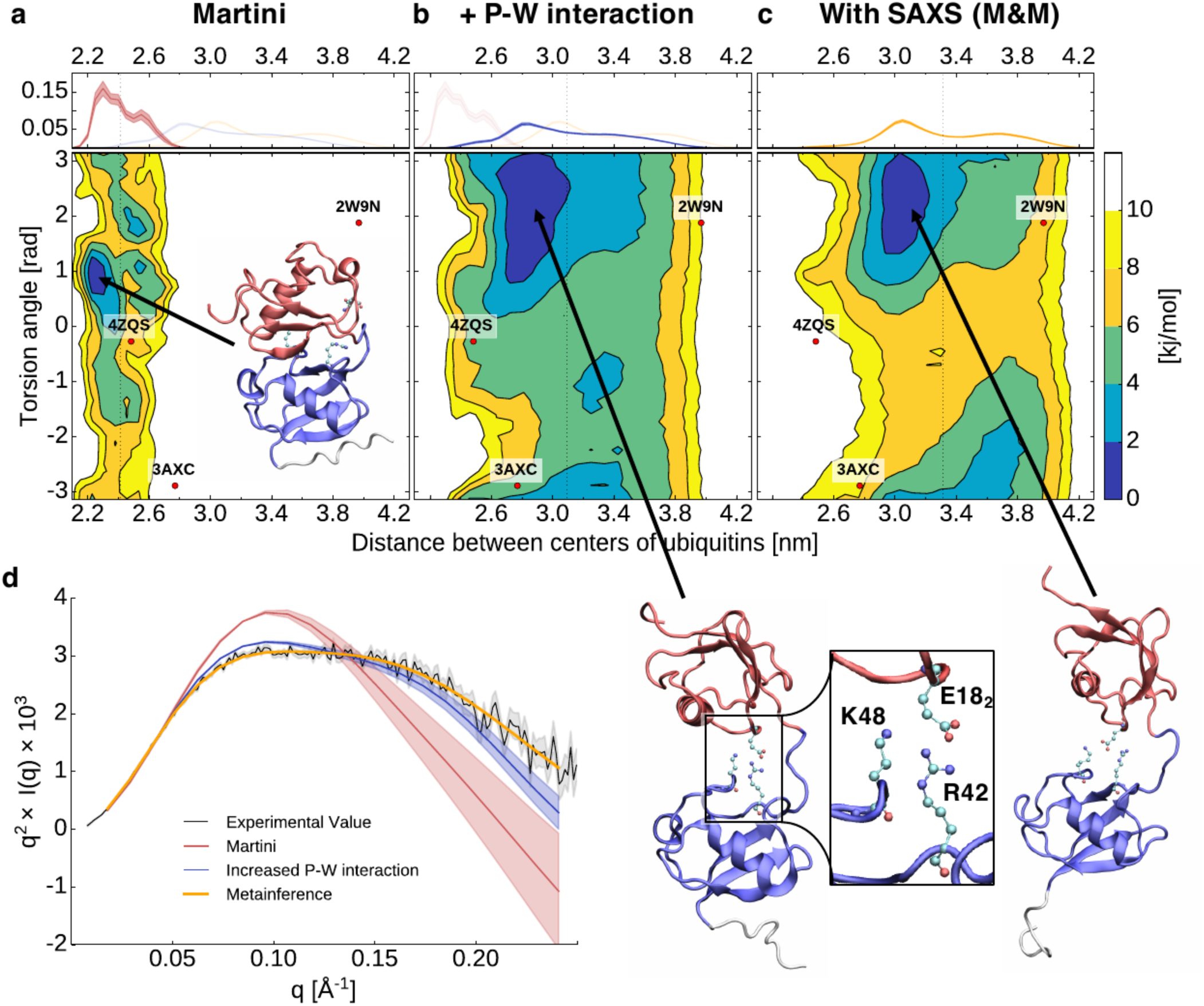
Characterization of the dynamics of linear di-ubiquitin. **a)-c)** Free energy landscapes (in kJ/mol) as a function of the distance between the center-of-mass of the two ubiquitin domains and their relative orientation (measured as the torsion angle between two axes defined using the first and second half of the sequence of each ubiquitin, see Methods). The dots represent the coordinates associated with the available di-ubiquitin crystal structures. On top is shown the probability distribution of the distance between the centers of the two ubiquitin domains. **d)** Experimental and from simulation calculated Kratky plot. The shaded area represents the error range.

### Metainference SAXS simulations of Martini di-ubiquitin quantitatively reproduce the experimental data

Metadynamic metainference (M&M)^45^ simulation (see Methods) for Martini linear Ub_2_ (including our modified water) result in an ensemble of configurations characterized by a flatter and broader free energy landscape (**Figure 2c**) and in quantitative agreement with the experimental SAXS (**Figure 2d**,**e**; **Fig. S1**). With respect to the unrestrained simulation, the average distance between the two domains increased from 3.10 ± 0.02 nm to 3.32 ± 0.02 nm. The radius of gyration of the ensemble of 2.23 ± 0.01 nm quantitatively agrees with that derived from SAXS of 2.23 ± 0.02 nm. Qualitatively the topology of the free energy landscape is comparable to the unrestrained simulation but translated to larger relative distances. Overall the free energy landscape is quite flat with relatively limited free energy differences indicating that the two ubiquitin-domain are relatively free to move with respect to each other. Therefore, Ub_2_ shows highly dynamical behavior, which cannot be described by a few individual structures. Instead, a full ensemble is required in agreement with previous findings on linear as well as other di-ubiquitins.

From the performance point of view, the SAXS on-the-fly calculation used by Metainference is computationally demanding, but the use of a coarse-grained representation makes it far more affordable with respect to the same simulation performed at full atomistic resolution (**Table S2**). The loss of performance resulting from the use of SAXS is justified by the increased accuracy of the resulting simulations. Note, that it is not required to calculate the Metainference SAXS restraint at every step of the simulation. Indeed, by calculating it every 5 steps, we obtained a quantitatively equivalent ensemble (**Fig. S2**) at a fraction of the computational cost (**Table S2**). Notably, using Metainference allows us also to sample the scaling value, which is necessary to compare the experimental and computed SAXS curves. For Ub_2_ we observed a 3% higher scaling value for the simulation with increased protein water interaction and a 9% higher scaling value just with the Martini force field compared to the Metainference solution (**Figure S1, Table S1**).

### Linear poly-ubiquitin chains are preferentially extended, do not show long-range correlations and can be described as self-avoiding polymers

To investigate the dynamics of linear Ub_3_ and Ub_4_, we performed SAXS experiments on both proteins at different concentrations (**Fig. S3**). The measured SAXS data were then employed to perform M&M simulations (cf. Methods, **Table S3**). Additionally, unrestrained simulations based only on Martini with our modified water were also performed. In **Figure 3** (see also **Fig. S1**), we show the comparison of the back-calculated SAXS with respect to the experimental measures of Ub_3_ and Ub_4_. The effect of our improved water diminishes for the longer poly-ubiquitin chains. A comparison of the radius of gyration for the Ub_2_, Ub_3_, and Ub_4_ ensembles show that while the unrestrained and restrained simulations sample a comparable range of compactness, the restrained simulations are shifted towards a more extended conformational space. The trend of the average radius of gyration (2.0, 2.7, and 3.3 nm for Ub_2_, Ub_3_, and Ub_4_, respectively, cf. Methods, **Table S2**) suggests an almost linear increase of the size of the protein with the number of ubiquitin monomers. The analysis of the free-energy landscape for the Ub_2_ couples in Ub_3_ and Ub_4_ (**Fig. S4**) shows qualitatively the same behavior, suggesting that the interdomain interactions are essentially only those between neighbor domains (i.e. between Ub_2_). For Ub_3_ and Ub_4_ this is confirmed by analyzing the free energy landscape of non-neighbor ubiquitin domains. Overall the free energy landscape is flatter for larger poly-ubiquitin chains indicating that the interaction with neighboring ubiquitin becomes less and less specific. Also, the distance (centers of the two ubiquitin domains) distribution shifts from a bimodal distribution for Ub_2_ to a flatter one for Ub_3_ and Ub_4_. We also observe that Ub_3_ samples more extended conformations for large distances > 4.0 nm, while both Ub_4_ and Ub_3_ are forming more compact conformation below 2.6 nm. In **Fig. S5a-c** the free energy profiles are shown for the first-and-third ubiquitin Ub_3_(1-3) in Ub_3_ as well as for the first-and-third Ub_4_(1-3), second-and-fourth Ub_4_(2-4). These landscapes are all qualitatively similar showing that the interaction between two non-neighbor ubiquitins is quite rare. The average distance between a 1-3 or 2-4 ubiquitins pair is around 6 nm, with an average angle of around 140°. The first ubiquitin does not influence the relative orientation of the third ubiquitin. The first and fourth Ub_4_(1-4) ubiquitin couple, as shown in **Fig. S5d**, behaves similarly. Interaction between the first and fourth ubiquitin are also rare. In most cases, the distance between both ubiquitins is around 8.5 nm. There is also no strong preference for a specific torsion angle between all four ubiquitins.

**Figure 3.**
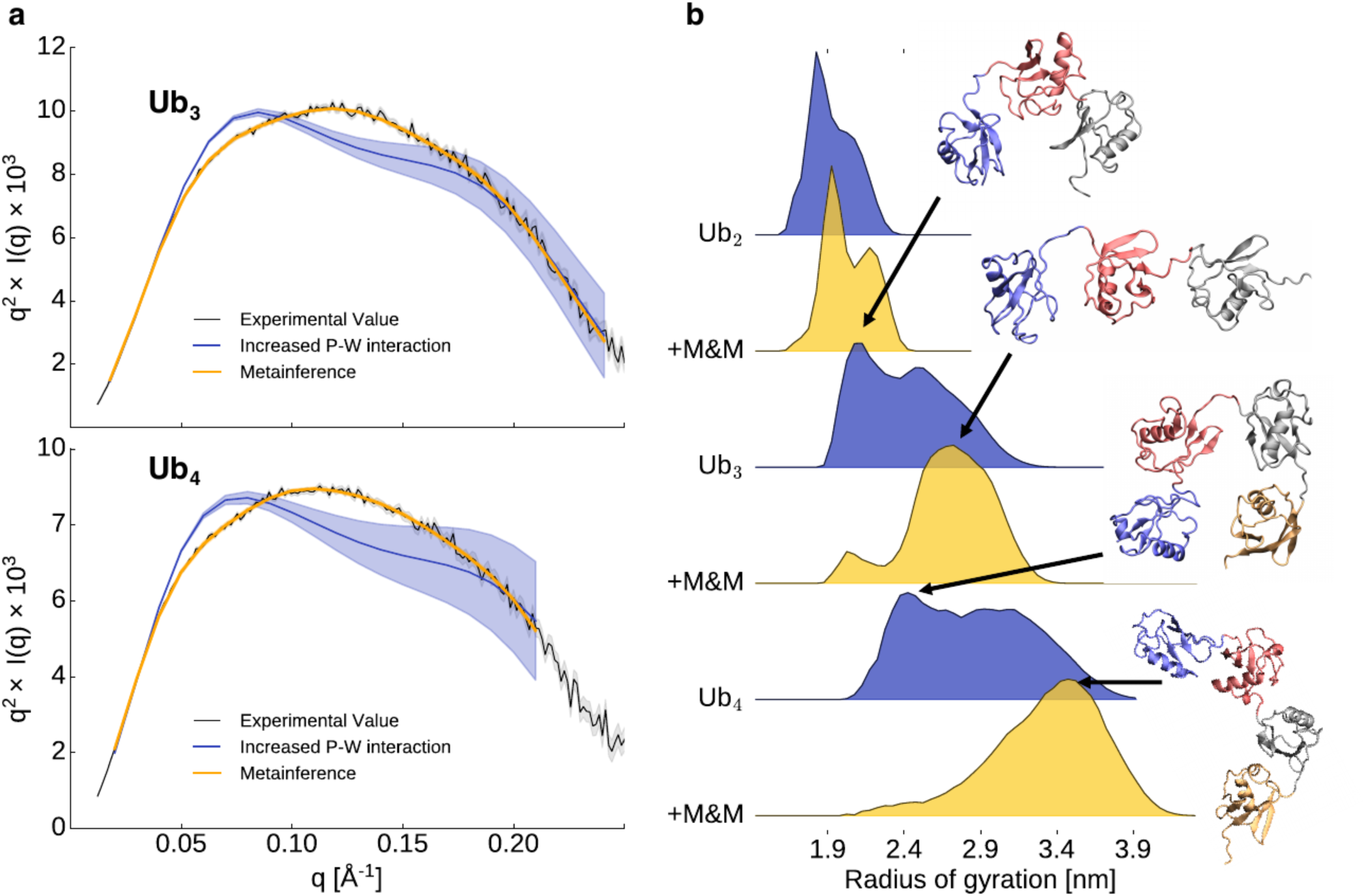
Characterization of the dynamics of linear tri- and tetraubiquitin. **a)** Experimental and from simulations calculated Kratky plot for tri- and tetraubiquitin. The shaded area represents the error range. **b)** Distribution of the radius of gyration from the ensemble with and without M&M.

To further assess the presence of short and long-range interactions between neighbor and non-neighbor ubiquitin couples we estimated the fraction of compact configurations by analyzing the minimum distance between neighbor and non-neighbor ubiquitin couples (**Figure 4a,b**). For neighbor and non-neighbor couples, there is a peak in the distribution around 0.5 nm. As already indicated by the free energy profiles, compact neighbor ubiquitin pairs represent around 40% to 50% of the ensemble, while contacts between non-neighbor couples are only present in around 8% for Ub_3_ and around 2% for Ub_4_, indicating an overall lack of compact states in linear poly-ubiquitins. A contact analysis for the Ub_2_ compact state indicates that this state is not structurally homogeneous. Even the most frequent contact is only present in 10 to 30% of all compact conformations, depending on the specific ubiquitin pair (**Figure 4c**). On the other hand, even the 10^th^ most frequent contact has still a probability between 5 to 15% while the 100th most frequent one is still in the 1 to 5% range.

**Figure 4.**
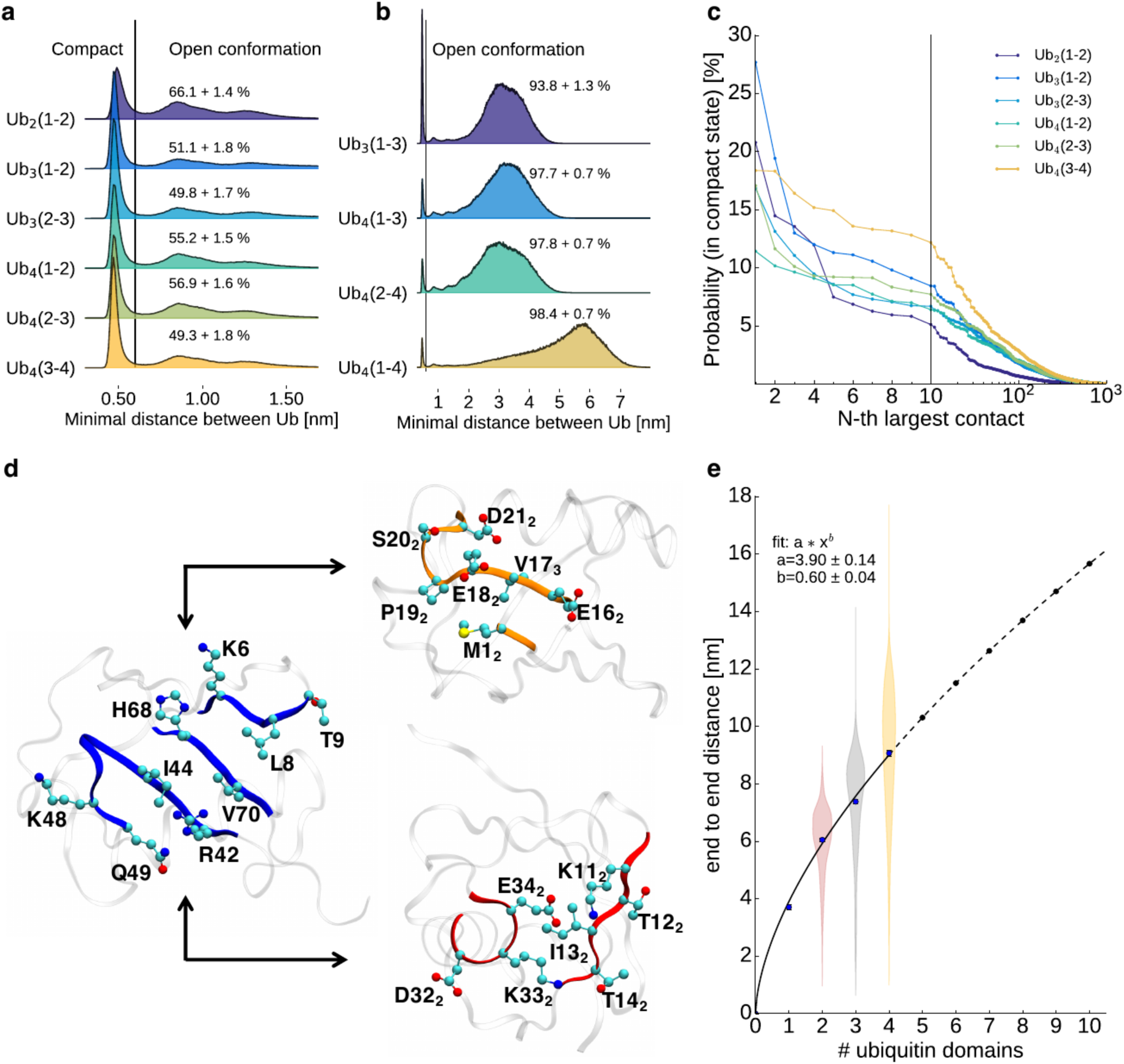
Intramolecular interactions of poly-ubiquitin. a) Minimum distance distribution between two neighboring ubiquitin cores (residue 1 to 70, residue 77 to 146, and so forth). Structures with a distance larger than 0.6 are defined as open. b) Minimum distance distribution between two non-neighboring ubiquitin cores. Structures with a distance larger than 0.6 are defined as open. c) The probability of finding contacts between two amino acids of neighboring ubiquitin cores d) Interaction surface of two neighboring ubiquitins. Residues from the blue marked surface (first ubiquitin - left) are interacting with residues of the orange marked surface (middle) or red marked surface (right) of the second ubiquitin. e) Average end to end distance of a linear poly-ubiquitin chain.

Nonetheless, all residues involved in the most frequent contacts belong to three distinct surfaces. These interactions define the preferred orientations between two adjacent ubiquitin pairs. All residues involved in the most frequent contacts from the first ubiquitin are on the same surface as the hydrophobic patch I44 (**Figure 4d, Fig. S6**), which is known to be also essential for interactions of ubiquitin with other proteins. The I44 surface interacts either with the surface around E18_2_ or I13_2_ (E18 or I13 would be the analog residues of the first ubiquitin). The E18_2_ surface is located opposite to the I44 surface while the I13_2_ surface is roughly 90° rotated to the I44 and E18_2_ surface. In **Fig. S6**, the ten most frequent contacts between ubiquitin cores for all adjacent ubiquitin pairs are illustrated. Ub_2_ is predominantly stabilized by salt bridges between the positive charged R42 and K48, and the negative charged E16_2_ and E18_2_. However, going to Ub_3_ and Ub_4_, electrostatic interactions become less important compared to Van der Waals (VdW) interactions (**Table S4**). For Ub_2_, the Coulomb interaction between charged amino acids is responsible for 27 % of the total interaction energy between two ubiquitin cores. This value goes as low as 9 % for the Ub(3-4) pair of Ub_4_. The lower electrostatic interactions may also explain the flatter free energy surfaces of Ub_3_ and Ub_4_ neighbor pairs (**Fig. S4**). On the other hand, the increased role of VdW interactions in Ub_3_ and Ub_4_ is compatible with the increase in the compact population of di-ubiquitin couples. Finally, while in Ub_2_ the I44 surface prefers to interact with the E18_2_ surface, interactions between the I44 and the I13_2_ surface are more important for the last pair of Ub_4,_ causing a shift of the preferred orientation between both ubiquitins.

Overall, our linear Ub_2_, Ub_3_, and Ub_4_ ensembles indicate that linear poly-ubiquitins are extended polymers, whose dynamics are mostly uncorrelated over a distance of more than one ubiquitin domain. In **Figure 4e** this behavior is further highlighted by plotting the end-to-end distance as a function of the number *N* of ubiquitin domain, *e*2*e*(*N*). Remarkably, fitting the data with a power law (including the end-to-end distance for Ub_1_) resulted in *e*2*e*(*N*) = 3.9*N*^0.60^ in remarkable agreement with Flory theory for self-avoiding polymers^46^. This is remarkable, since generally, proteins do not behave as self-avoiding chain, showing also less entropy in the denatured state. It is tempting to speculate that all poly-ubiquitins may be described as self-avoiding polymers following the same relationship for the end-to-end distance but with a different pre-factor (i.e., characteristic length) associated with the distance between the C-terminus glycine and the specific linkage side chain. For K63-linked poly-ubiquitins, we could test this, even if only with the mono-ubiquitin and di-ubiquitin, making use of a SAXS based ensemble we recently published^47^. By fixing the exponent to 0.6 and setting the prefactor to the average distance between the C-terminus and K63, the expected *e2e* distance of Ub_2_ fits the ensemble. This suggests that the fit can be used to predict the behavior for longer K63-linked Ub_n_ chains (**Fig. S7)**. Extrapolating for other linkages we observe that K11- and K48-linked poly-ubiquitin, which are known to populate more compact states in solution^48-49^, have a shorter distance between the C-terminus and the lysine and would consequently show smaller pre-factors and populate systematically more compact states than K63- and M1-linked.

### The conformational entropy of long linear poly-ubiquitins modulates NEMO binding

In order to study the dynamics and changes of linear poly-ubiquitin dynamics upon binding to cognate proteins, we used simulations and experiments to characterize the interaction of the NEMO UBAN domain (NEMO_258–350_) to the linear poly-ubiquitins Ub_2_, Ub_3_ and Ub_4_. The NEMO UBAN domain is a dimer in solution^23^. Previous studies have shown two different binding stoichiometries in solution and crystalline state for Ub_2_: with either two NEMO monomers bound to one Ub_2_(2:1) or two NEMO monomers bound to two Ub_2_ (2:2)^23-24^. Poly-ubiquitin chains longer than Ub_2_ harbor potentially more than one binding site and could thus bind more than one NEMO dimer with a theoretical stoichiometry for Ub_N_ of 2(N-1):1 (NEMO:Ub_N_). A crystal structure (PDB 5H07) shows the binding of two linear Ub_3_ to four ABIN monomers (a homologous of NEMO which also form a dimer in solution)^50^. The observed binding mode requires Ub_3_ to be in a relatively compact and univocally oriented configuration in order to avoid steric hindrances between the two ABIN dimers. This would dramatically decrease the entropy not only of each di-ubiquitin couple but also that of the overall chain and thus should be entropically disfavored in solution (**Fig. S5**). Indeed, isothermal titration calorimetry (ITC) experiments show that in solution only one ABIN2 dimer binds to Ub_3_^50^, arguing that higher stoichiometries are artifactual, induced by crystal packing and do not reflect the solution assembly.

To characterize the binding in solution of NEMO to Ub_3_ and Ub_4_ we performed SAXS (**Figure 5a,b, Fig. S3, Table S5**), ITC (**Figure 5c,d, Fig. S8, Table 1, Table S6**), Size Exclusion Chromatography (SEC) coupled with static light scattering (SLS) (**Fig. S9, Table 2, Table S7**), and surface plasmon resonance (SPR) (**Fig. S10, Table S8**). Extending Ub_2_ to Ub_3_ and Ub_4_ does not affect the (2:1) binding stoichiometry, with either poly-ubiquitin protein binding two NEMO monomers (one NEMO dimer) (**Figure 5, Table 2**). In SEC experiments with SLS (**Table S7**) we detected NEMO:Ub_3_ complexes with molecular weights ranging from (41-45 kDa), which is similar to the MW of the calculated 2:1 NEMO:Ub_3_ complex (47.7 kDa), while for NEMO:Ub_4_ complexes only a single peak is found with MW between 53-56 kDa (calculated MW 2:1 NEMO:Ub_4_ 56.2 kDa) (**Table 2, Fig. S9, Table S5**). SAXS measurements (**Figure 5a-b, Table 2, Fig.S3**) confirmed the stoichiometry observed by SLS-SEC. ITC (**Figure 5c,d, Fig. S8, Table 1, Table S6**) indicates that NEMO binds to Ub_3_ and Ub_4_ with very similar enthalpy (ΔH of -17.9 and -18.8 kJ/mol, respectively) suggesting that the molecular interactions and binding interfaces between NEMO and the different poly-ubiquitins are similar to the one described for NEMO:Ub_2_^23^ (ΔH of -16.9 kJ/mol). Affinities for Ub_3_ and Ub_4_ are 1.6 and 4.1 µM close to 1.8 µM obtained for Ub_2._ ITC data comparing shorter and longer poly-ubiquitins may suggest that longer poly-ubiquitins can form long range, flanking interactions with NEMO, resulting in a gain of enthalpy and loss of entropy with respect to shorter ones. SPR confirmed the binding between NEMO and Ub_3_ and Ub_4_ with equilibrium dissociation constants (*K*_D_) of 9.6 µM and 6.4 µM for Ub_3_ and Ub_4,_ respectively (**Fig. S10, Table S8**).

**Table 1.** Isothermal titration calorimetry (ITC) measurement of the NEMO interaction with Ub_2_, Ub_3,_ and Ub_4_. NEMO was titrated into the ubiquitin solutions in 50 mM sodium phosphate pH 7, 50 mM NaCl. Values are averages ± standard errors from three measurements. The individual ITC curves are shown in Fig. S8. ^*^ Experiments taken from Vincendeau et al^23^. A stoichiometry of N=2 corresponds to one NEMO dimer binding to one poly-ubiquitin protein.

| NEMO + | Ub<sub>2</sub>* | Ub<sub>3</sub> | Ub<sub>4</sub> |
| --- | --- | --- | --- |
| **N** | 1.86 ± 0.01 | 1.84 ± 0.09 | 2.02 ± 0.04 |
| **K<sub>D</sub> [μM]** | 1.81 ± 0.12 | 1.57 ± 0.28 | 4.13 ± 0.30 |
| **ΔH [kJ/mol]** | -16.9 ± 0.1 | -17.9 ± 0.6 | -18.8 ± 0.1 |
| **-TΔS [kJ/mol]** | -15.8 | -15.3 ± 0.4 | -11.9 ± 0.2 |

**Table 2.** Molecular weight determination. SAXS and size exclusion chromatography (SEC) in combination with static light scattering (SLS) were used to determine the molecular weight of NEMO, Ub_3_, Ub_4_, NEMO:Ub_3_ and NEMO:Ub_4_. The conditions were 50 mM Tris.HCl pH 8, 300 mM NaCl.

| Protein | MW<sub>exp</sub> [kDa]<br>(SAXS) | MW<sub>exp</sub> [kDa]<br>(SEC/SLS) | MW<sub>calc</sub><br>[kDa] |
| --- | --- | --- | --- |
| **NEMO dimer** | 24.4 | 20.6 | 22.0 |
| **Ub<sub>3</sub>** | 20.5 | 24.1 | 25.7 |
| **Ub<sub>4</sub>** | 29.9 | 32.3 | 34.2 |
| **NEMO:Ub<sub>3</sub> 2:1** |  | 41.1 | 47.7 |
| **NEMO:Ub<sub>4</sub> 2:1** |  | 56.2 | 56.2 |
| **NEMO:Ub<sub>3</sub> (1.4:1)** | 40.0 |  | 40.8 |
| **NEMO:Ub<sub>3</sub> (2.7:1)** | 38.7 |  | 40.6 |
| **NEMO:Ub<sub>4</sub> (1.0:1)** | 37.9 |  | 46.6 |
| **NEMO:Ub<sub>4</sub> (3.1:1)** | 45.1 |  | 44.0 |

**Figure 5.**
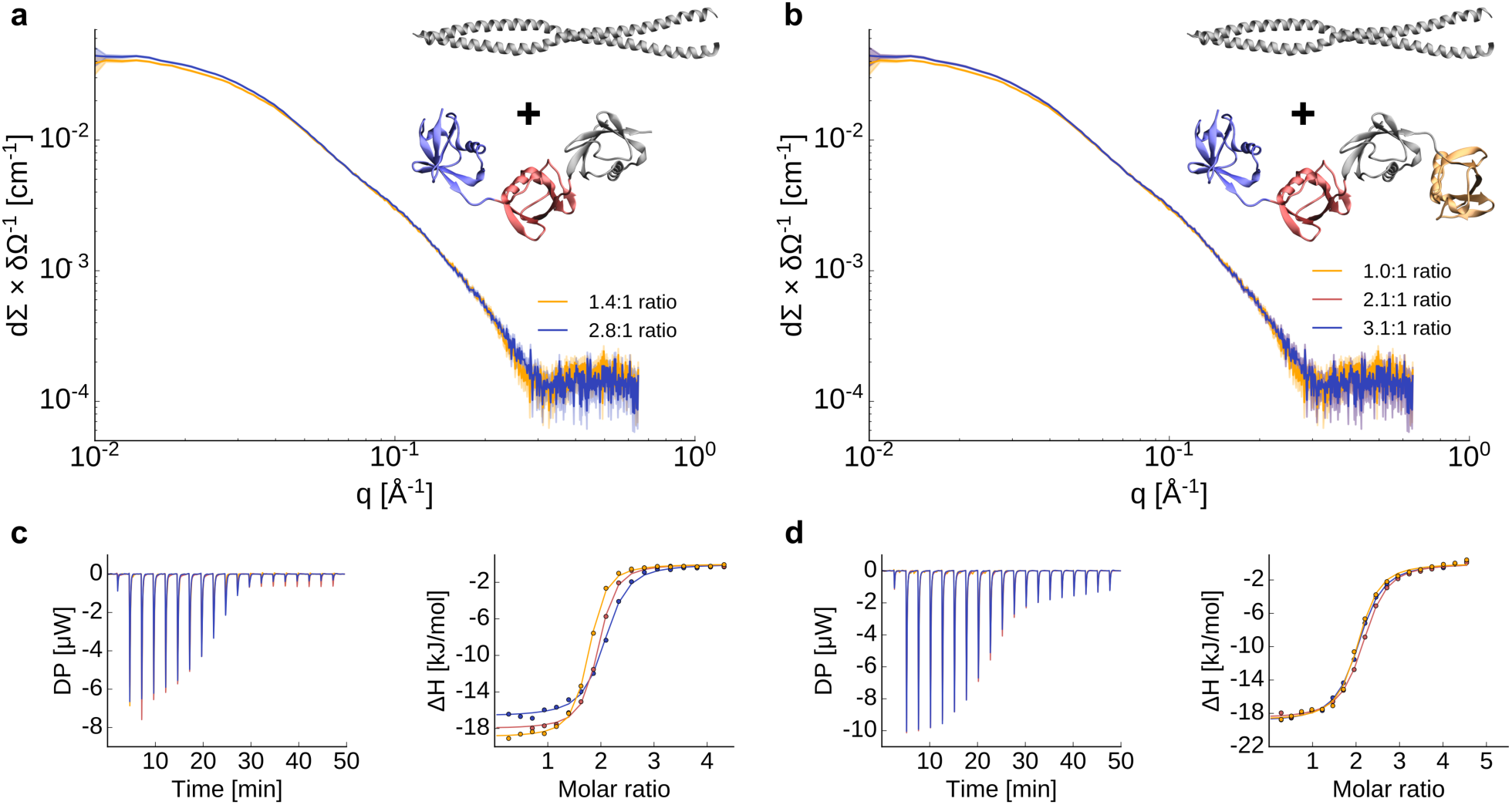
Effect of chain length on the binding of NEMO. **a)** and **b)** SAXS experiments for different rations of NEMO and Ub_3_ (a) and Ub_4_ (b). **c)** and **d)** Isothermal titration calorimetry (ITC) measurement of the interaction of NEMO with -Ub_3_ (c) and Ub_4_ (d). NEMO was titrated into the poly-ubiquitin solutions. The experiment was repeated three times.

To explain the contradictory observation of the stoichiometry in solution and in crystal and to better understand the molecular recognition between linear poly-ubiquitins and NEMO, we characterized the dynamics of a NEMO_258–350_:Ub_2_ complex. An M&M Martini simulation was performed including SAXS data for the complex previously measured (**Figure 6a, Fig. S11**). The resulting ensemble of structures highlights how binding to NEMO strongly decreases the conformational freedom of linear Ub_2_ (**Figure 6a, Figure 2**). Neither the ensemble nor the crystal structures of other bound Ub_2_ are located close to the minima of the free Ub_2_ ensemble, which has a different distance and orientation between both ubiquitins. Interestingly, Ub_2_ residues building up the NEMO_258–350_:Ub_2_ interface overlap with those involved in the interdomain interactions (**Fig. S11**), in particular residues around the hydrophobic patch I44 and the previously mentioned E18_2_ surface. The observed interaction sites are in agreement with observed NMR chemical shifts perturbations reported in Vincendeau et al^23^. The ensemble also provides a possible explanation for the different binding stoichiometry observed in solution (NEMO:Ub_2_ 2:1) and crystalline state (NEMO:Ub_2_ 2:2). A detailed analysis of the NEMO binding sites indicates that almost all the residues of the NEMO unoccupied site are less exposed to the solvent than those on the occupied one (**Fig. S12**). While the solvent accessible surface area (SASA) of the occupied site is 13 nm^2^ the one of the unoccupied is 11.9 nm^2^ showing that both binding sites are not equal after the binding of Ub_2_. Altogether these observations indicate that, in solution, binding of linear Ub_2_ to NEMO_UBAN_ induces allosteric effects that modulate the overall structure and dynamics of the NEMO dimer. These observations also suggest that the 2:2 highly symmetric binding mode observed in the dense and ordered crystalline state becomes entropically unfavorable to the more flexible and far less dense solution state of the complex.

**Figure 6.**
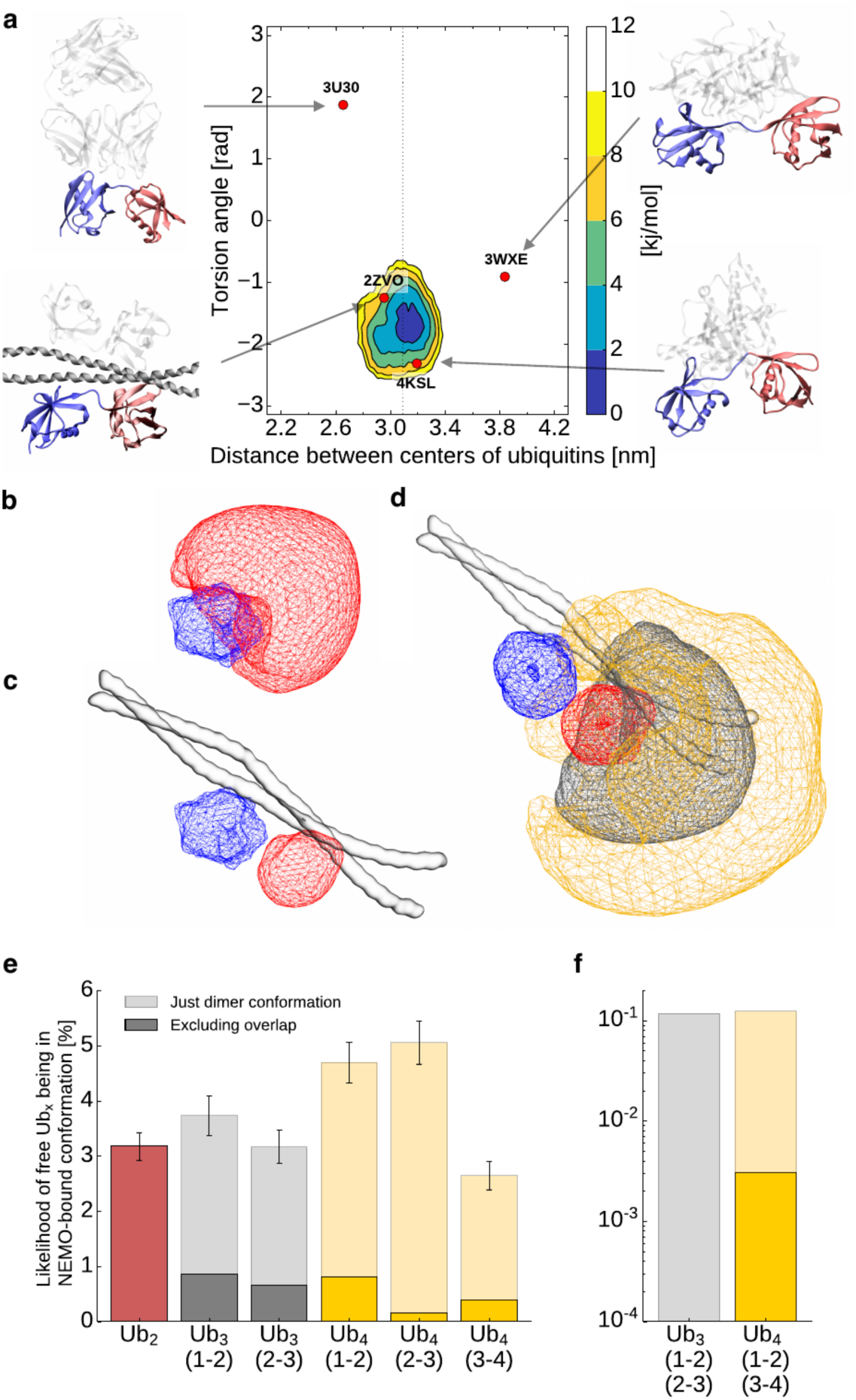
Comparison between free and NEMO-bound poly-ubiquitin ensembles. **a)** Free energy landscapes (in kJ/mol) as a function of the distance between the centers of the two ubiquitin domains and their relative orientation for Ub_2_ bound to NEMO. The dots represent the coordinates associated with the available crystal structures with Ub_2_ bound to different proteins. **b-c)** Conformational space of free (b) and NEMO-bound (c) ubiquitin pairs in Ub_2_. The blue area represents the first ubiquitin, while the red area shows the conformation of the second ubiquitin relative to the first one. **d)** Conformational space of third (gray area) and forth (orange area) ubiquitin in Ub_4_ with the first Ub-pair being in a NEMO-bound conformation. **e-f)** Probability of free Ub_2_, Ub_3,_ and Ub_4_ of being in a NEMO-bound conformation for one (e) or two (f) NEMO dimers. The transparent bars show the likelihood of the individual pairs being in the NEMO-bound conformation (RMSD < 6 A compared to the average Ub_2_ structure in the NEMO-bound simulation). The dark bars show the probability of being in the NEMO-bound conformation, excluding structures with an overlap between the non-bound ubiquitins and NEMO.

## Discussion

Structural biology investigations on poly-ubiquitins have mostly focused on di-ubiquitins observing that different protein linkages correspond to different protein dynamics leading to different exposed regions for the binding with partners^47-49, 51-56^. Ubiquitin signaling has been found associated not only to the linkage type but also to the length of the ubiquitin chains^8-10^. Here we first develop an efficient and accurate integrative approach to characterize the conformational ensembles of linear poly-ubiquitin by combining the Martini coarse-grain force field with SAXS experiments in the framework of Metainference. We then employ our method to try to rationalize the length dependent behavior of linear poly-ubiquitins and the consequence for the interaction with their partner NEMO. **Figure 6** rationalizes the observed differences in binding by comparing our free Ub_N_ ensembles with our NEMO-bound ensemble. The fraction of bound-like configurations in the Ub_2_ ensemble is a small fraction of the total ensemble suggesting a large conformational entropy loss upon binding. This is likely compensated by a release of a large number of water molecules from the binding interfaces upon binding in order to result in a final entropy gain as indicated by ITC (**Table 1**). The probability of finding at least one di-ubiquitin pair in a bound-like configuration in the Ub_3_ and Ub_4_ ensemble increases slightly more than linearly (3.2%, 6.7%, and 12.0% for Ub_2_, Ub_3_ and Ub_4_, respectively) suggesting that longer poly-ubiquitins are likely to bind NEMO more favorably with respect to shorter ones. In fact, the SEC-SLS experiments show that NEMO bound to Ub_4_ eluted as a single bound peak in comparison with Ub_3_ (**Table S7** and **Fig. S9**). Since both NEMO:Ub_4_ and NEMO:Ub_3_ have similar *K*_D_s this can indicate a difference in kinetic stability. To provide a structural interpretation for this hyphothesis, we calculated the actual probability of finding the full poly-ubiquitin in a configuration compatible with the binding (and thus avoiding configurations that would lead to a steric clash with NEMO (**Figure 6b-d**)) from our free polyubiquitin ensembles. The probability decreases from Ub_2_ to Ub_4_ (3.2%, 1.5% and 1.4% for Ub_2_, Ub_3,_ and Ub_4_, respectively) which can lead to entropy loss. At the same time, non-specific flanking interactions between poly-ubiquitin and NEMO far from the binding site can increase the enthalpy. This is also in agreement with previous measures where, using a longer NEMO construct (NEMO_242-419_) that could provide more surface for interactions, affinities of 3 and 0.3 µM were reported for Ub_2_ and Ub_4_, respectively^57^. This principle is also common for intrinsically disordered proteins possibly modulating the lifetime of complexes^58^. Such long-range effects would be less pronounced for a less entropic chain and can play a length dependent role in the overall interaction.

Interestingly, the probability of finding two di-ubiquitin pairs in a bound like configuration is essentially negligible for both Ub_3_ and Ub_4_ providing a rationale why a 2:1 NEMO_258–350_:Ub_4_ interaction is favored by entropy with respect to the 4:1. Making use of our polymer model, we can also speculate that a long-enough poly-ubiquitin may be able to bind two NEMO dimers with a higher order (i.e. 4:1) stoichiometry with respect to the 2:1 observed for the Ub_2_-Ub_4_ range. Given that the end-to-end distance of Ub_2_ corresponds to one half of that of our NEMO construct (5.8 nm for Ub_2_ and 11.4 nm for NEMO), and that the two Ub_2_ units that bind the two NEMO should be allowed to be flexible, one can estimate the length of this poly-ubiquitin to be such that *e*2*e*(*N* − 2) ≥ 11.4nm. This results in a minimum length of *N=8* ubiquitins. While this result will not be quantitative when considering a full-length NEMO, it suggests a possible need for such long chains in the assembly of the IKK complex.

## Conclusions

The combined use of experiments and molecular dynamics simulation is a powerful tool to investigate the structure and dynamics of biomolecules and provide a ground for the functional interpretation of protein dynamics. Here, we combined SAXS and coarse-grained Martini simulations to study the conformational dynamics of linear poly-ubiquitins and their binding to NEMO accurately and efficiently. The resulting conformational ensembles allowed us to propose that linear-poly-ubiquitin behave as self-avoiding polymer chains. This might also apply for poly-ubiquitins in general (with a different characteristic lengths). Combining structural studies with multiple biophysical experiments, we provide a systematic assessment of the effect of the poly-ubiquitin chain length in the molecular recognition of cognate proteins, suggesting that poly-ubiquitin may modulate the binding with their partners in a length-dependent manner.

## Methods

### Coarse-grained molecular dynamics simulations

Coarse-grained molecular dynamics simulations (CG-MD) were applied to investigate the dynamic of linear di-tri- and tetraubiquitin (Ub_2_, Ub_3_, Ub_4_), as well as Ub_2_ with bound NEMO. In total, 11 different simulations have been performed with a total simulation length of 780 µs. An overview of all simulations can be found in **Table S3**.

All CG-simulations were run using Gromacs 2016.3^59^ and the Martini force field^34-35^. Additionally, an elastic network model was used to conserve the secondary and tertiary structure. In the case of poly-ubiquitin, the elastic network inside was only defined for the core region (from residue 1 to 70, 77 to 146, 153 to 222 or 229 to 298) and not between different domains nor for the linker region. All simulations were performed with periodic boundary conditions, and the systems were solvated with a 0.1 M NaCl solution and run as an isothermal-isobaric (NpT) with a temperature of 300 K and a pressure of 1 bar using 20 fs time steps. To control the temperature and pressure, the v-rescale thermostat^60^ was used with a coupling constant τ_t_ = 1.0 ps together with the Parrinello-Rahman barostat^61^ and a coupling constant of τ_p_ = 20.0 ps and compressibility of χ = 3.0 × 10^−4^ bar^−1^. Nonbonded interactions were treated with a dielectric constant of 15 and using a cutoff distance of 1.1 nm. VMD was used for visualization^62^.

For many simulations, the Martini 2.2. force field was modified to increase the protein-water (P-W) interaction, which was achieved by giving water beads their own atom type with a 5% larger C6 parameter in interactions with all other atom types resulting in around 5% higher protein-water interactions.

Parallel-biased metadynamics^37, 63^, as implemented in PLUMED2^64^, was utilized to enhance the sampling of the conformational space, together with the multiple walker approach^65^ with 112 replicas for free poly-ubiquitin or 64 replicas for free NEMO as well as bound NEMO with Ub_2_, where each replica had a different starting conformation. The employed collective variables were the distances between the centers of the different ubiquitin cores, a torsion angle between the centers of residue 1 to 36 and 37 to 70 of two different ubiquitins, the radius of gyration (calculated only with backbone atoms) and the alphabeta collective variable describing the torsional angles for linkers between the ubiquitin pairs. In total, four collective variables for Ub_2_, 9 for Ub_3_ and 16 for Ub_4_ were used. In the case of simulations with NEMO, additional collective variables were employed for the shape of NEMO and distances between Ub_2_ and NEMO. The bias factor of the well-tempered metadynamics^66^ was set to 10, the frequency for the hill addition was 200 (every 4 ps), the height of gaussian hills was 0.1 kJ/mol for simulations with di- and Ub_3_, 0.075 kJ/mol for simulations with tetraubiquitin and 0.02 kJ/mol for the simulation with NEMO bound to Ub_2_. The flexible Gaussian approach^67^ was used to determine the Gaussian width during the simulation.

Metainference^36^, a method based on Bayesian inference, was used to integrate experimental SAXS data into simulations and was coupled with metadynamics (M&M)^45^. The calculation of the SAXS intensities from a CG Martini representation is implemented in the PLUMED-ISDB module^68-69^ using the parameters derived by Niebling et al^70^ and the Debye equation. The SAXS data of the different systems were fitted with a 16^th^-degree polynomial to calculate points used for restraints. 21 equidistant points between for q between 0.017 to 0.24 nm^-1^ were used for Ub_2_, Ub_3_ and for free and bound NEMO, 19 points for q between 0.025 to 0.19 nm^-1^ for Ub_4_. The rage depends on the quality of the experimental data. An initial scaling value was determent by comparing the calculated and experimental SAXS intensities for the lowest q value. Metainference was used with the outlier noise model^36^ for each data point, and the restraints were applied every 5^th^ step. A scaling faction and offset for the experimental data were sampled using a flat prior between 0.9 to 1.1 or -1 to 1. The error for calculating an average quantity σ_mean_ was determined automatically^71^ from the maximum standard error over 2 ps of simulations.

Six different simulations with Ub_2_ were performed using: Martini 2.2 with metadynamics, with increased P-W-interaction and metadynamics, with M&M, with M&M applied every step, and with Martini 3 beta. Notably, Martini 3 beta was not stable with metadynamics, therefor 112 replicas were run on a longer time scale. The SAXS data of Ub_2_ were taken from Vincendeau et al^23^. Notably, the profile was measured with Ub_2_ containing a HIS-tag on the N-terminus which was also modeled. Ub_3_ and Ub_4_ simulations were run with increased P-W interaction, metadynamics with and without Metainference and SAXS. All poly-ubiquitin simulations were run for at least 500 ns per replica. The simulations with free NEMO and NEMO bound to Ub_2_ were performed with increased P-W interaction and M&M with SAXS for at least 100 ns per replica. The SAXS data of NEMO bound to Ub_2_ were taken from Vincendeau et al^23^.

The plumed input files as well as the modified Martini topology files are deposited in PLUMED-NEST^72^ as plumID:20.009.

### Protein expression and purification

Human NEMO_258–350_ C347S was expressed and purified as described in Vincendeau et al^23^. Protein concentration was determined by measuring the absorbance at 205 nm using specific absorbance for NEMO_258–350_ C347S of 300990 M^-1^ cm^-1^, respectively^73^.

The constructs for the expression of Ub_3_ and Ub_4_ were a kind gift of Dr. Paul Elliott and Dr. David Komander (MRC Laboratory of Molecular Biology, Cambridge, UK). The constructs were transformed into *E. coli* strain BL21 (DE3) and cultured at 20°C in 2-L flasks containing 500 ml ZYM 5052 auto-induction medium^74^ and 100 µg/ml carbeniicillin. Cells were harvested by centrifugation after reaching saturation, resuspended in 60 ml lysis buffer (50mM Tris-HCl, 300mM NaCl, 10mM MgCl_2_, 10 µg/ml DNaseI, 1mM AEBSF.HCl, 0.2% (v/v) NP-40, 1 mg/ml lysozyme, pH 8.0), and lysed by sonication. The lysate was clarified by centrifugation (40,000 x *g*) and filtration (0.2 µM). The supernatant was heated in a water bath for 10-15 minutes at 60°C and the precipitate removed by centrifugation. The supernatant was dialyzed overnight against 2 L of buffer A (50 mM sodium acetate pH 4.5), clarified by centrifugation and applied to a 5-ml HiTrap SP HP column (GE Healthcare), equilibrated in buffer A. Bound proteins were eluted using a linear gradient (10 column volumes) from 0 to 1 M NaCl in buffer A using an Äkta Purifier (GE Healthcare). Elution fractions (1.6 ml) were collected in wells containing 250 µl 1M Tris-HCl pH 9.0. Fractions containing Ub_3_ or Ub_4_ were pooled, concentrated and applied to a HiLoad 16/600 Superdex 75 column (GE Healthcare), equilibrated in buffer B (50mM Tris-HCl, 100mM NaCl, pH 7.4). The main elution peak containing Ub_3_ or Ub_4_ was collected and concentrated to approx. 3-6 mg/ml, flash frozen and stored at -80°C. Protein concentrations were determined by measuring the absorbance at 205 nm using specific absorbance for Ub_3_ and Ub_4_ of 747790 and 997980 M^-1^ cm^-1^, respectively^73^. The Ub_4_ concentration values used on interaction studies of Ub_4_ with NEMO were corrected by 30% based on the SEC/SLS results.

### Small angle X-ray scattering measurements

SAXS measurements were performed on a Rigaku BIOSAXS1000 instrument with an HF007 microfocus generator equipped with a Cu-target at 40 kV and 30mA. Transmissions were measured with a photodiode beamstop, q-calibration was made by an Ag-behenate measurement. Absolute calibration was done with calibrated glassy carbon^75^. Measurements were done in four 900 second frames, which were averaged. Under these conditions, no radiation damage was detected. Circular averaging and background subtraction were done with the Rigaku SAXSLab software v 3.0.1r1.

Radii of gyration were calculated with the ATSAS package v 2.8.0^76^. Fits for the MW determination were made in Origin v 9. SAXS measurements were made at 293 K using a buffer contained 300mM NaCl and 50mM Tris-HCl at a pH of 8.0. Experiments on the free proteins were performed at the following concentrations (**Table S5**): NEMO at 2.34, 4.62 and 7.72 mg/ml, Ub_3_ at 3.41, 6.72 and 11.17 mg/ml, Ub_4_ at 4.5, 9.1 and 15.1 mg/ml. Experiments with NEMO and Ub_3_ and Ub_4_ at different ratios were performed at two concentrations (between 3-4 and 7-8 mg/ml) at the following ratios: NEMO:Ub_3_ at 1.4:1 and 2.7:1 ratios, NEMO:Ub_4_ at 1.0:1, 2.1:1 and 3.1:1 ratios. No concentration-dependent effects were detected.

### Isothermal Titration Calorimetry

ITC measurements were carried out at 298 K using a PEAQ-ITC titration microcalorimeter (MicroCal, Malvern). The NEMO to Ub_3_ calorimetric titration consisted of 19 injections of 2 μl of a 2.13 mM NEMO solution, into the reaction cell containing 300 μl of 94.71 μM Ub_3_, at a stirring speed of 750 rpm. The NEMO to Ub_4_ calorimetric titration consisted of 19 injections of 2 μl of a 2.84 mM NEMO solution, into the reaction cell containing 300 μl of 120.6 μM Ub_4_, at a stirring speed of 750 rpm. Sample conditions were 50 mM sodium phosphate pH 7.0 and 50 mM NaCl. The heat of dilution was obtained by titrating NEMO into the sample cell containing only buffer. Experiments were done in triplicate. The ITC data were analyzed using the software MicroCal PEAQ-ITC analysis software. Parameters are presented as averages ± standard errors.

### Size Exclusion Chromatography with Static Light Scattering

Static light scattering (SLS) experiments were performed of NEMO mutant (C347S) in complex with tri- and tetra-ubiquitin at 30°C using a Viscotek TDA 305 triple array detector (Malvern Instruments) downstream to an Äkta Purifier (GE Healthcare) equipped with an analytical size exclusion column (Superdex 200 10/300 GL, GE Healthcare) at 4°C. The samples were run at approx. 8 mg/ml at a flow rate of 0.5 ml/min. The experiments were performed using a Tris buffer (50 mM Tris-HCl, 300 mM NaCl, pH 8.0) and a phosphate buffer (50 mM sodium phosphate, 50 mM NaCl, pH 7.0). The molecular masses of the samples were calculated from the refractive index and right-angle light-scattering signals using Omnisec (Malvern Instruments). The SLS detector was calibrated with a 4 mg/ml BSA solution using 66.4 kDa for the BSA monomer and a *dn/dc* value of 0.185 ml/g for all protein samples.

### Surface Plasmon Resonance measurements

SPR measurements were performed at 25°C using a Pioneer FE instrument (FortéBio, Molecular Devices). Ub_3_ and Ub_4_ were covalently immobilized on to two different flow cell channels on a biosensor chip by amine coupling to 456 and 721 RU, respectively, using a 10 mM NaOAc pH 5 immobilization buffer. NEMO was injected in a two-fold concentration series over immobilized ubiquitins at 30 µL/min flow rate using a PBS running buffer (50 mM sodium phosphate, 50 mM NaCl, 0.005% tween, pH 7). The data were analyzed using Qdat Data Analysis Tool version 2.6.3.0 (FortéBio). The sensorgrams were corrected for buffer effects and unspecific binding to the chip matrix by subtraction of blank and reference surface (a blank flow cell channel activated by injection of EDC/NHS and inactivated by injection of ethanolamine). The equilibrium dissociation constants (K_D_) were estimated by plotting responses at equilibrium (Req) against the injected concentration and curve fitted to a Langmuir (1:1) binding isotherm.

## Acknowledgments

The authors acknowledge Paul Elliott and David Komander (MRC Laboratory of Molecular Biology, Cambridge, UK) for the Ub_3_ and Ub_4_ plasmids. We acknowledge Daniel Krappmann (Helmholtz Zentrum München, Germany) for reading the paper and providing useful feedback. AJ and CC acknowledge support by the Technische Universität München-Institute for Advanced Study, funded by the German Excellence Initiative and the European Union Seventh Framework Programme under Grant Agreement No. 291763. SMØS. MS acknowledges funding by the DFG SFB1035, and AB acknowledge the support by the Lundbeck Foundation (Grant R190-2014-3710). We gratefully acknowledge the Gauss Centre for Supercomputing e.V. (www.gauss-centre.eu) for funding this project by providing computing time on the GCS Super-computer SuperMUC at the Leibniz Supercomputing Center (LRZ, www.lrz.de). We acknowledge SAXS measurements at the facility of the SFB1035 at Department Chemie, Technische Universität München.

