## Supporting Information for "Molecular recognition and dynamics of linear poly-ubiquitins: integrating coarse-grain simulations and experiments"

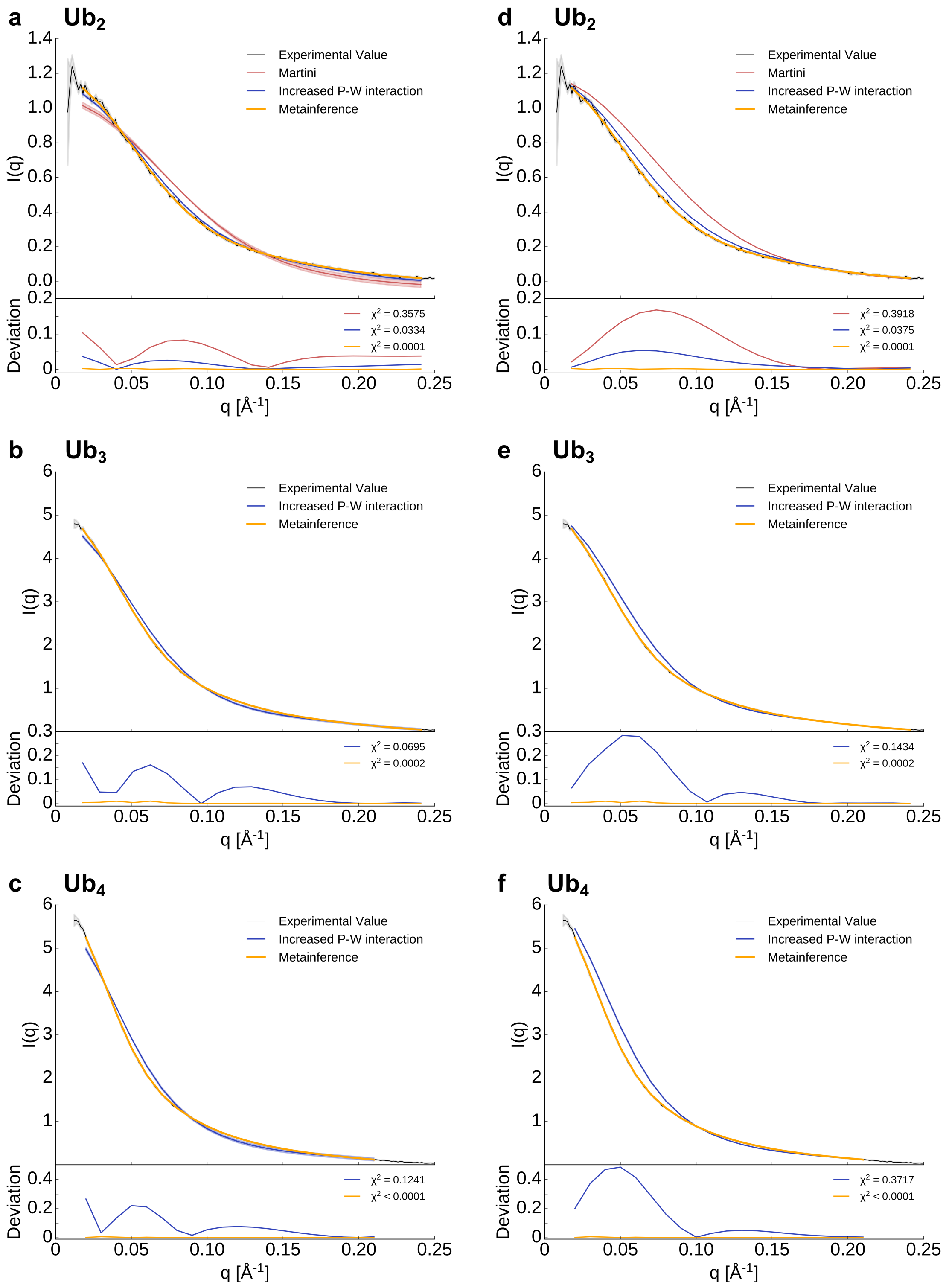

**Fig. S1.** Experimental and theoretical SAXS curves of polyubiquitin**.** a-c) SAXS intensities obtained from simulations fitted to the experimental curves of $Ub_2$, $Ub_3$, and $Ub_4$. d-f) SAXS intensities obtained from simulation with the scaling factor obtained by the M&M simulation. The scaling value should be similar for all structures and ensembles of the same protein since the SAXS intensities for scattering vectors close to q = 0 $Å^{-1}$ are mostly independent of the structure and dynamics of the system. In many cases, the SAXS curve calculated directly from a given structure or ensemble is simply fitted to the experimental data (a-c). This approach produces scaling values that are generally not transferable between different ensembles. Using the same scaling value (d-f) leads to a reduction of the agreement between theoretical and experimental SAXS profiles for non-Metainference ensembles.

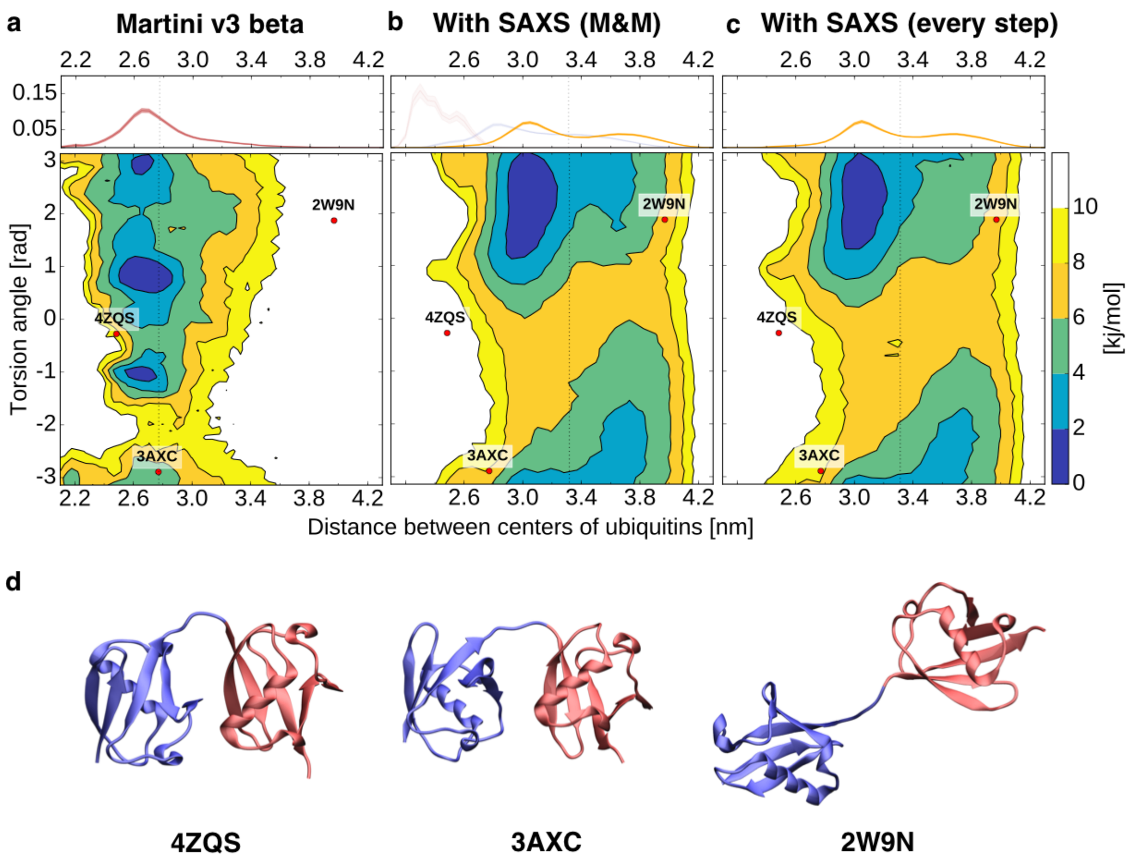

**Fig. S2. Free energy surfaces of linear di-ubiquitin for different simulation setups.** a)-c) Free energy landscapes (in kJ/mol) as a function of the distance between the centers of the two ubiquitin domains and their relative orientation. The dots represent the coordinates associated with the available di-ubiquitin crystal structures. On top is shown the probability distribution of the distance between the centers of the two ubiquitin domains. d) Crystal structures of free $Ub_2$.

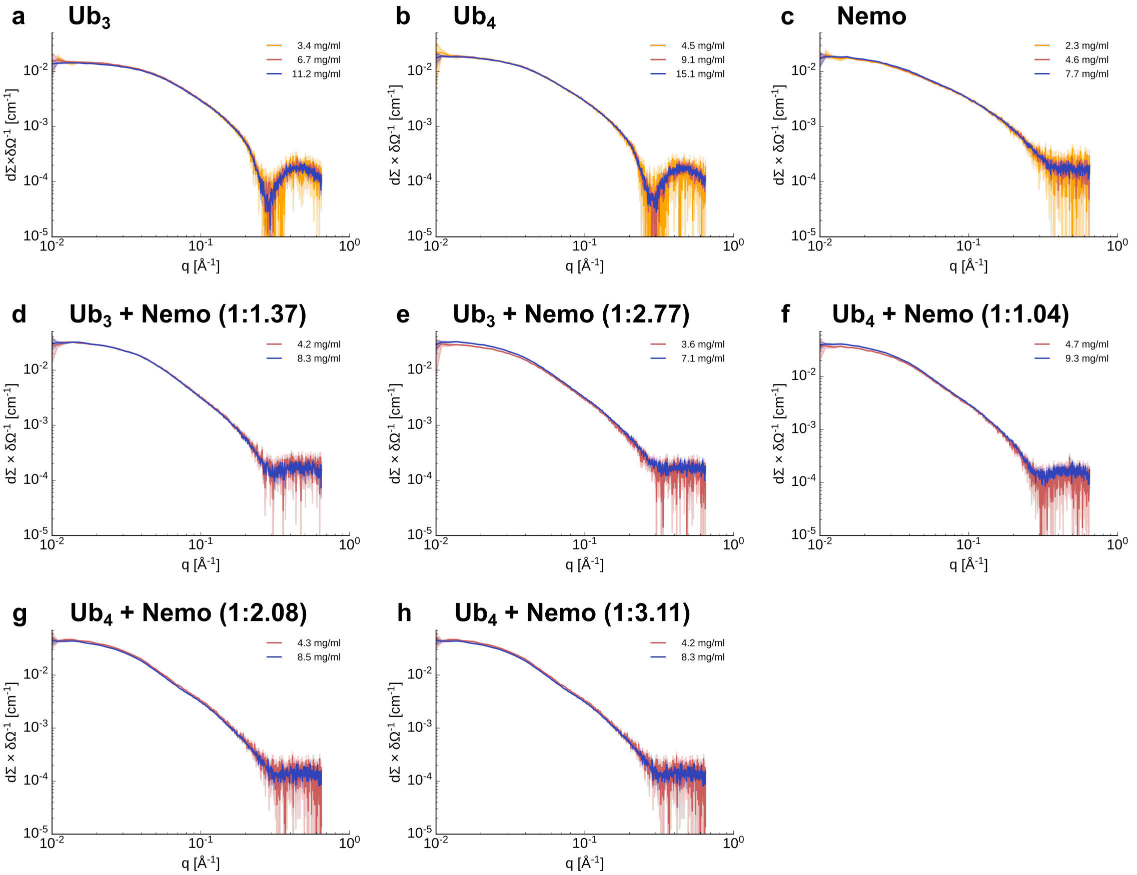

**Fig. S3. Experimental SAXS curves with different protein concentration** a) Linear $Ub_3$ b) linear $Ub_4$ c) NEMO d) $Ub_3$ with NEMO in a 1:1.37 ratio e) $Ub_3$ with NEMO in a 1:2.77 ratio f) $Ub_4$ with NEMO in a 1:1.04 ratio g) $Ub_4$ with NEMO in a 1:2.08 ratio h) $Ub_4$ with NEMO in a 1:3.11 ratio

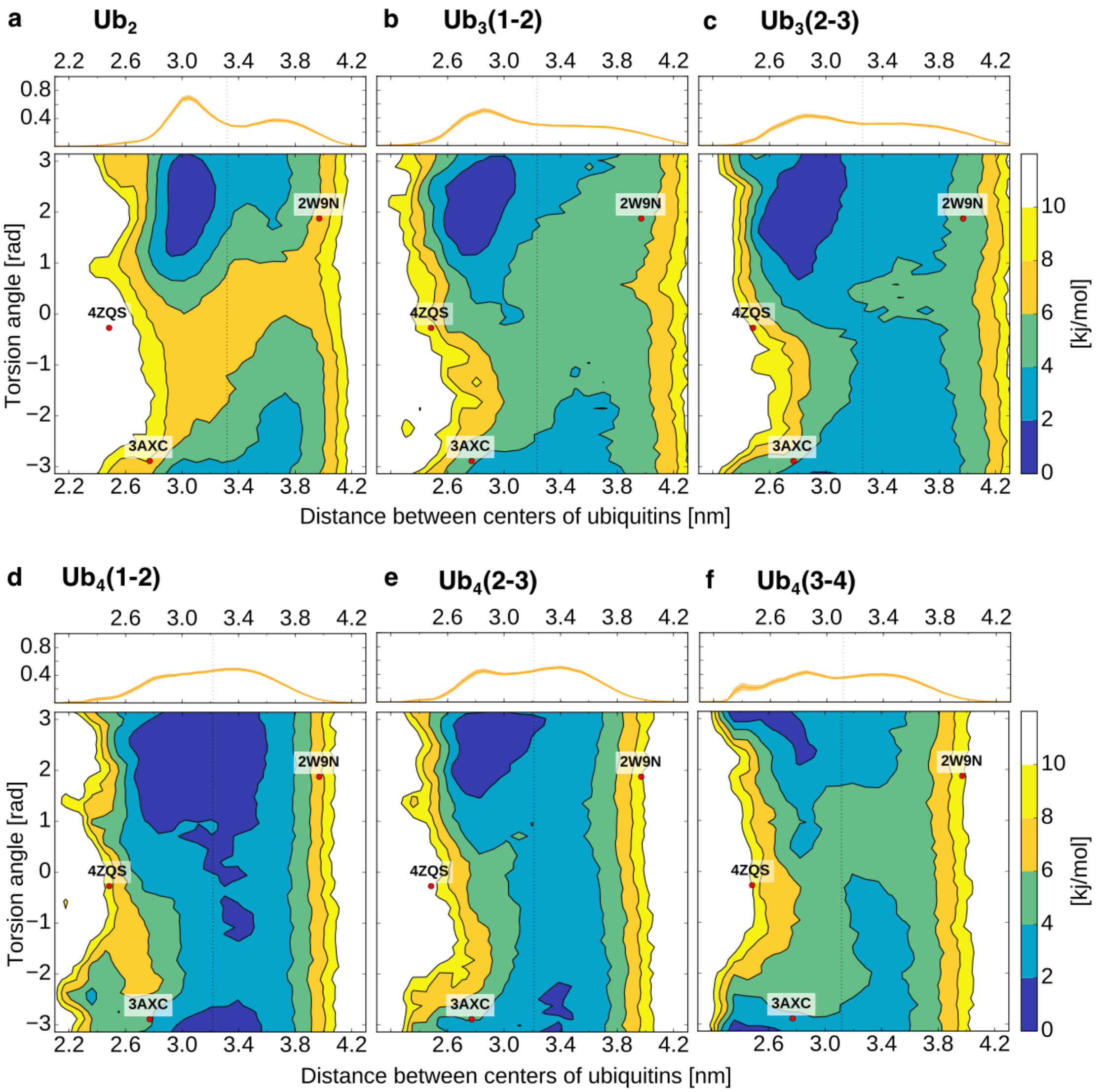

**Fig. S4. Free energy surfaces for all neighboring ubiquitin pairs of $Ub_2$, $Ub_3$, and $Ub_4$.** a)-f) Free energy landscapes (in kJ/mol) as a function of the distance between the centers of the two ubiquitin domains and their relative orientation. The dots represent the coordinates associated with the available di-ubiquitin crystal structures. On top is shown the probability distribution of the distance between the centers of the two ubiquitin domains.

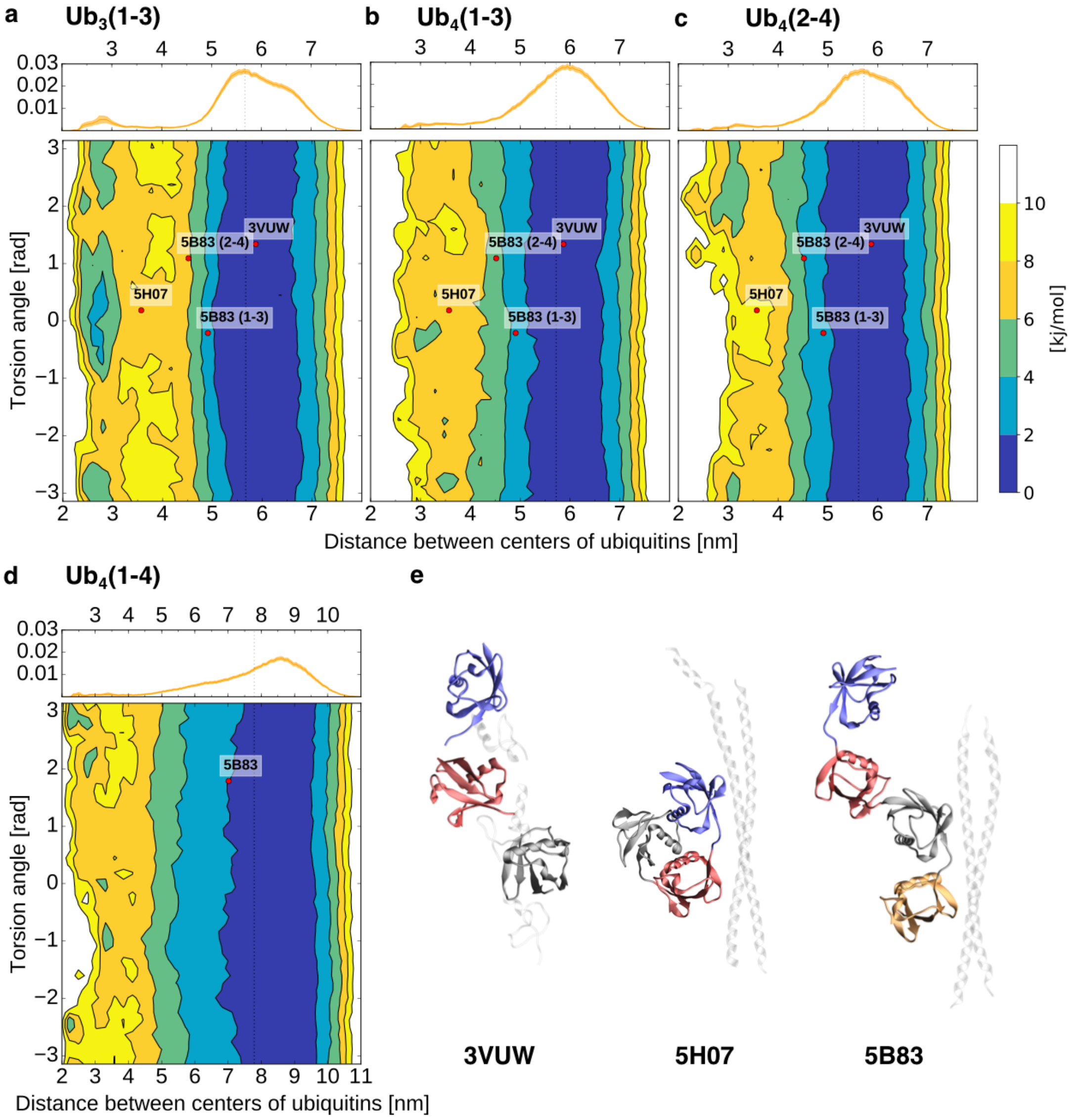

**Fig. S5. Free energy surfaces for all non-neighboring ubiquitin pairs of $Ub_3$ and $Ub_4$.** a)-d) Free energy landscapes (in kJ/mol) as a function of the distance between the centers of the two ubiquitin domains and their relative orientation. The dots represent the coordinates associated with the available di-ubiquitin crystal structures. On top is shown the probability distribution of the distance between the centers of the two ubiquitin domains. e) Crystal structures of linear $Ub_3$ and $Ub_4$ in a bound conformation (3VUW, 5H07, 5B83).

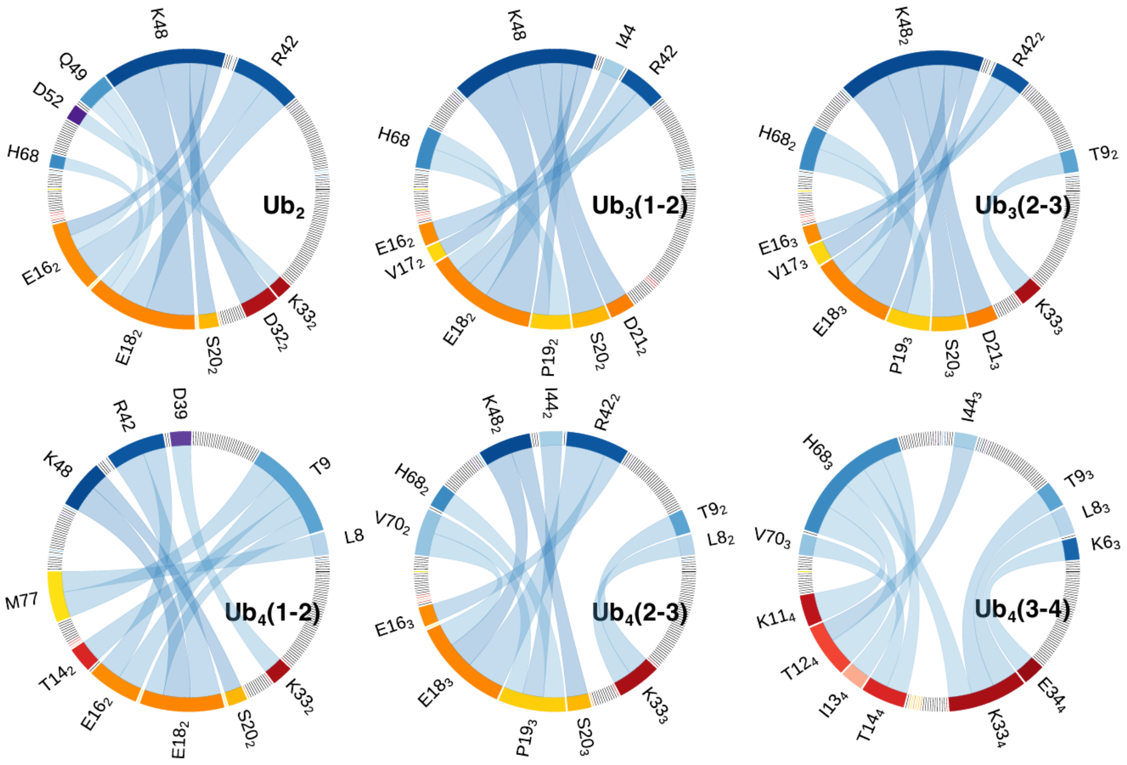

**Fig. S6. Chord diagram of the ten most common contacts between two close-by ubiquitins.** The edge colors are corresponding to the colors of the interaction surfaces shown in **Figure 4d**. The thickness of the edge-edge connection is correlated to the probability of finding this specific interaction (thicker = more likely). The transparency of the connections depends on the type of interaction Low, intermediate or high transparency is related to charged, polar or hydrophobic interactions.

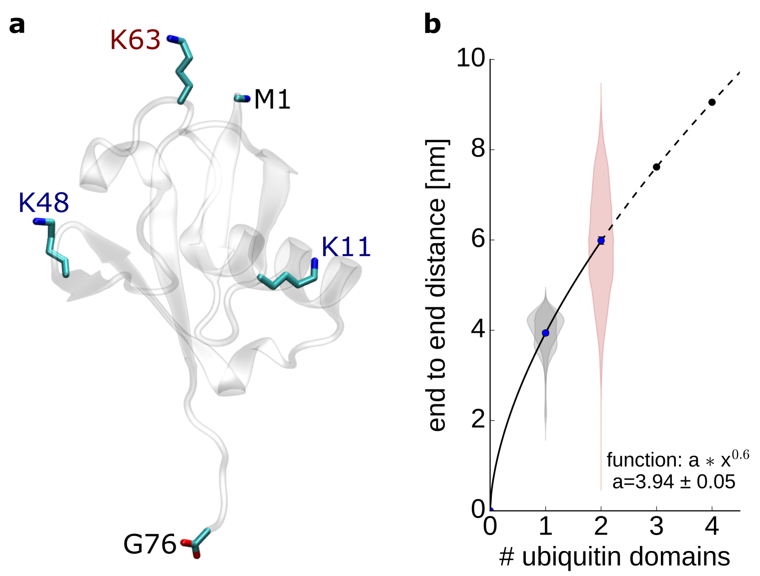

**Figure S7. End to end distances in ubiquitin. a)** Start and end residues in the ubiquitin crystal structure. K63 side chain is further away from G76 than M1 backbone. K48 and K11 are closer to G76. **b)** Average end to end (e2e) distance of K63 poly-ubiquitin chain. The end to end (e2e) distance of N=1 was determined as the average e2e distance of the proximal and distal ubiquitin in K63 diubiquitin.

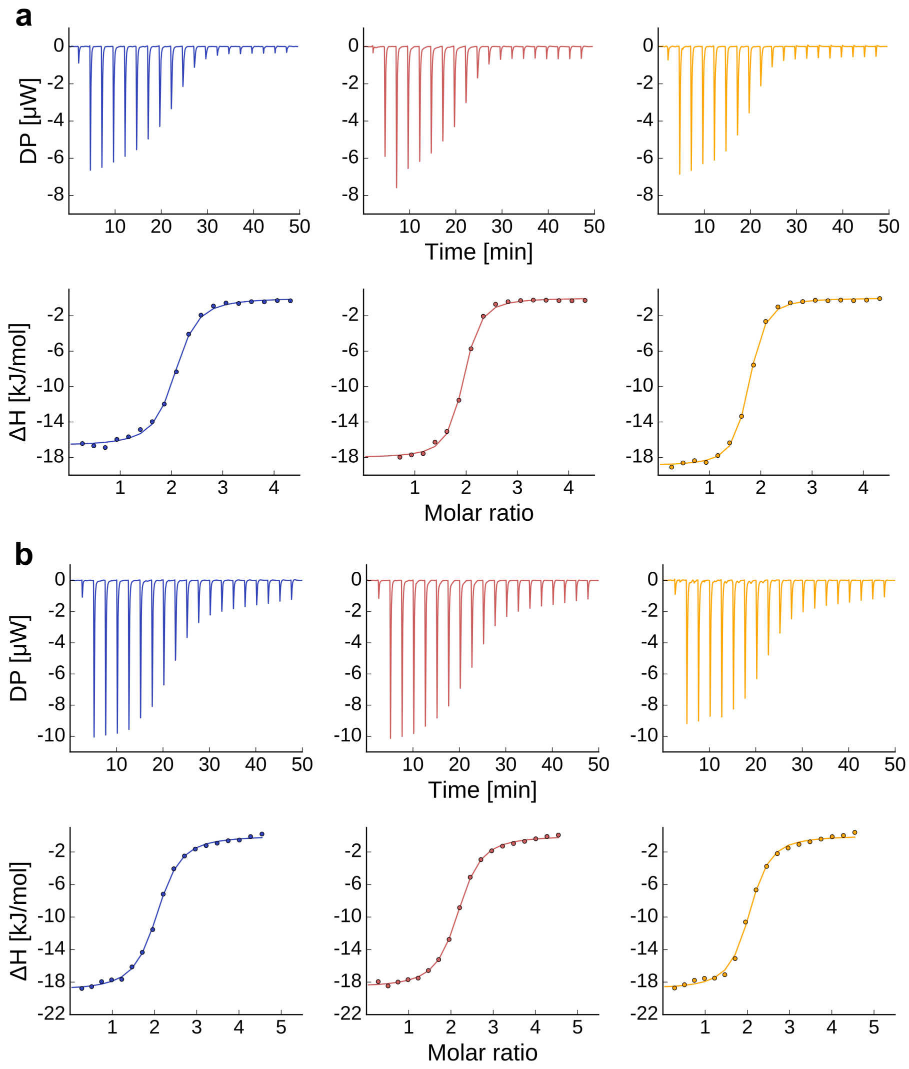

**Figure S8. Isothermal titration calorimetry (ITC) measurement of the interaction of NEMO with $Ub_3$ (a) and $Ub_4$ (b).** $NEMO_{258–350}$ was titrated into the polyubiquitin solutions.

**a** **b**

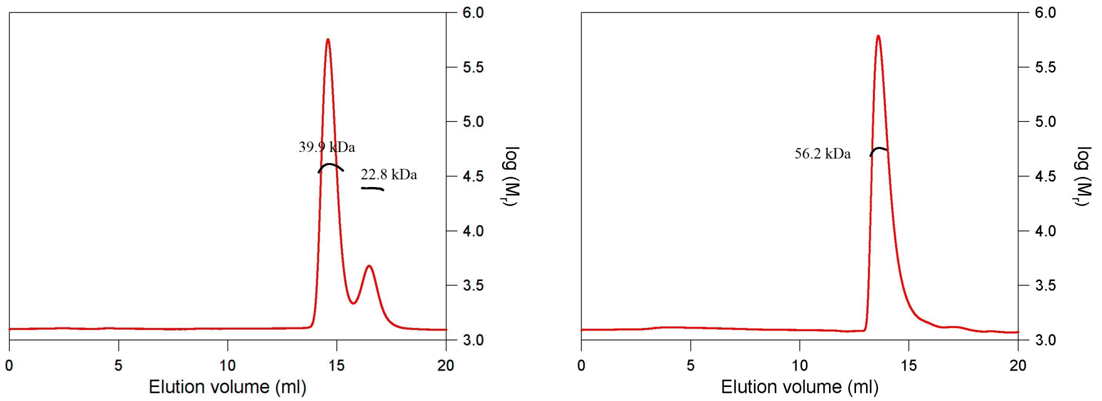

**Figure S9. Determination of the molecular weight of NEMO:Ub3 and NEMO:Ub4 complexes using size exclusion chromatography (SEC) in combination with static light scattering (SLS). a,b)** The refractive index (red) and right-angle light scattering (not shown) signals were monitored and used to determine the molecular weights (black). The complexes NEMO:Ub3 and NEMO:Ub4 were mixed in a 1.4:1 and a 2.3:1 (NEMO:Ub) molar ratio, respectively, and 100 µl of the samples (8 mg/ml) were applied to a Superdex 200 10/300 GL column. **a**) The NEMO-Ub3 sample elutes in two peaks. The first peak elutes at 14.6 ml and a molecular weight (MW) of 39.9 kDa. This peak contained a mixture of the NEMO:Ub3 complex and the remaining free NEMO dimer. The second peak elutes at 16.5 ml and contained the remaining Ub3 (MW of 24.6 kDa). **b**) The NEMO:Ub4 complex eluted in one major peak at 13.6 ml and a MW of 56.2 kDa indicating that indeed a 2:1 complex is formed.

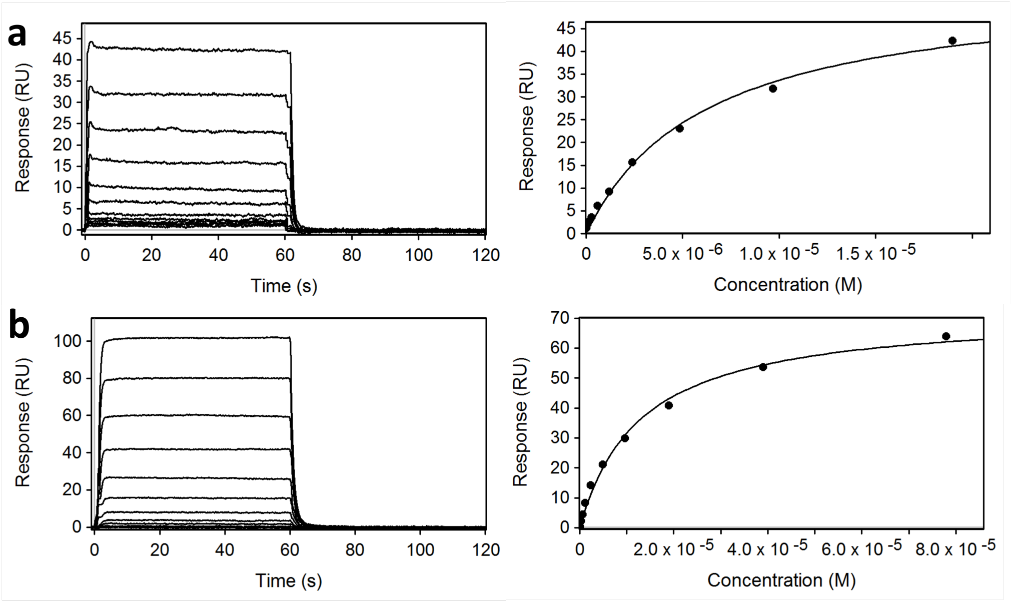

**Figure S10. SPR sensorgrams of NEMO interacting with immobilized $Ub_3$ or $Ub_4$.** NEMO was injected in two-fold serial dilution ranging from 0.9 –19 µM over immobilized $Ub_3$ (456 RU). **a)** and $Ub_4$ (721 RU). **b)** (Left panels). Sensorgrams are blank injection and reference surface subtracted. An activated and inactivated blank surface was used as a reference. Right panels display plots of equilibrium binding responses at the end of the analyte injections from sensorgrams in left panels against analyte concentration. Steady state equilibrium dissociation constants ($K_D$), from curves fitted to a 1:1 model, are presented in **Table S8**.

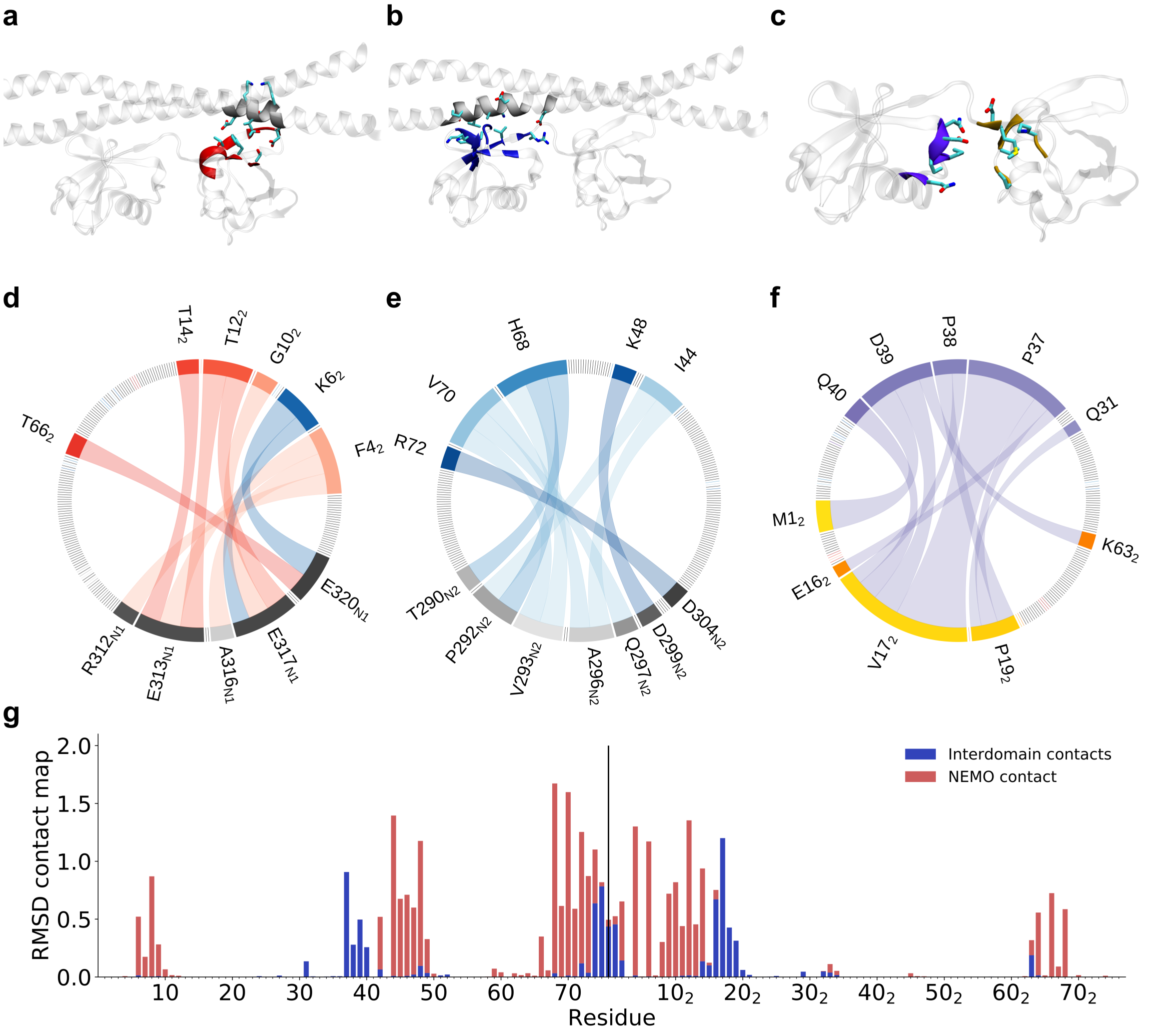

**Figure S11. Intramolecular interactions of di-ubiquitin and NEMO. a)** Interaction surfaces of Ub$_2$ and NEMO 1. The red surface is comparable to the red area marked in **Figure 4d**. **b)** Interaction surfaces of Ub$_2$ and NEMO 2. The blue surface is similar to the blue area marked in **Figure 4d**. **c)** Interaction surface of two neighboring ubiquitins. **d-f)** Chord diagram of the ten most common contacts between d) NEMO 1 and Ub$_2$ e) NEMO 2 and Ub$_2$ f) both ubiquitin cores. The edge colors are corresponding to the colors of the interaction surfaces shown in a-c. The thickness of the edge-edge connection is correlated to the probability of finding this specific interaction (thicker = more likely). The transparency of the connections depends on the type of interaction. Low, intermediate, or high transparency is related to charged, polar, or hydrophobic interactions. **g)** Contact map differences between free and NEMO-bound Ub$_2$.

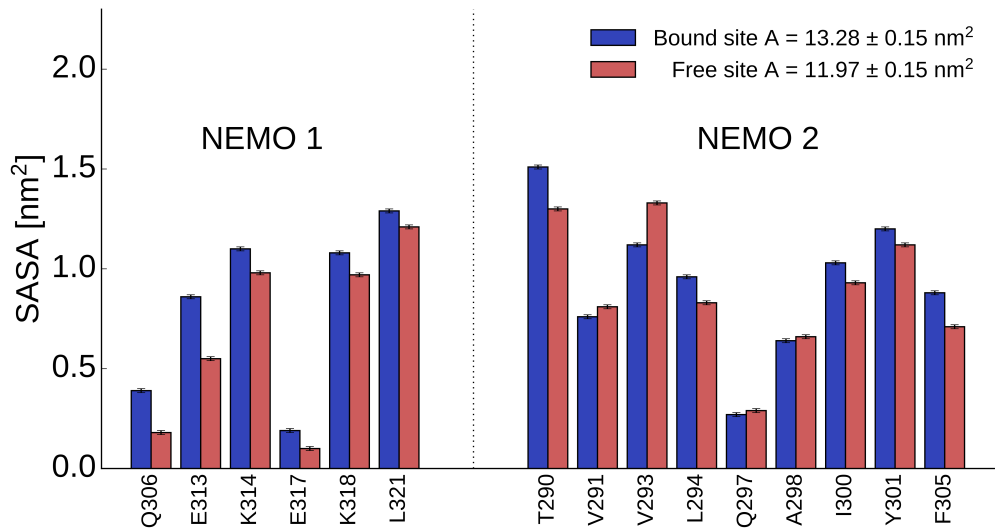

**Figure S12. Solvent accessible surface area (SASA) changes in NEMO dimer upon binding to $Ub_2$.** The bars show the SASA per residue, averaged over the conformational ensemble, for the residues involved in the binding with $Ub_2$. Errors estimated by block analysis. Blue bars are for the two NEMO monomers in the ubiquitin bound state, while the red bars are for the free state. On top is reported the SASA for all the considered residues.

**Table S1. Comparison of the quality of the SAXS calculations for different simulation setups.**

| | Scaling value | Offset | Chi-Square | $R_g$ | $R_g$ with expression tag |
|---|---|---|---|---|---|
| *Di-ubiquitin* | | | | | |
| **Martini** | 1.093 +- 0.037 | 0.036 +- 0.018 | 0.3575 | 1.654 +- 0.005 | 1.729 +- 0.004 |
| **Increased P-W interaction** | 1.034 +- 0.010 | 0.151 +- 0.005 | 0.0334 | 1.933 +- 0.006 | 2.050 +- 0.006 |
| **With SAXS** | 1.003 +- 0.001 | 0.005 +- 0.0004 | 0.0001 | 2.034 +- 0.006 | 2.230 +- 0.006 |
| | | *Experimental Value:* | | | 2.23 +- 0.02 |
| *Tri-ubiquitin* | | | | | |
| **Increased P-W interaction** | 1.042 +- 0.013 | 0.015 +- 0.023 | 0.0695 | 2.417 +- 0.013 | |
| **With SAXS** | 0.990 +- 0.001 | 0.021 +- 0.001 | 0.0002 | 2.699 +- 0.013 | |
| | | *Experimental Value:* | | 2.66 +- 0.02 | |
| *Tetra-ubiquitin* | | | | | |
| **Increased P-W interaction** | 1.079 +- 0.019 | 0.008 +- 0.037 | 0.1241 | 2.851 +- 0.022 | |
| **With SAXS** | 0.983 +- 0.0003 | 0.032 +- 0.001 | < 0.0001 | 3.332 +- 0.018 | |
| | | *Experimental Value:* | | 3.301 +- 0.04 | |

**Table S2. Performance for the different simulation setups. All values are in ns/day.**

| Cores per replica | 1 | 2 | 4 | 8 | 16 |
|---|---|---|---|---|---|
| *Di-ubiquitin* | | | | | |
| **Martini** | 415 | 730 | 1296 | 2098 | 3228 |
| **Martini + Metadynamics** | 352 | 564 | 928 | 1292 | 1590 |
| **With SAXS** | 142 | 255 | 466 | 740 | 1088 |
| **With SAXS - every step** | 40 | 78 | 149 | 254 | 412 |
| **Atomistic** | < 1 | 1 | 2 | 3 | 6 |
| *Tri-ubiquitin* | | | | | |
| **With SAXS** | 98 | 182 | 329 | 541 | 816 |
| *Tetra-ubiquitin* | | | | | |
| **With SAXS** | 51 | 95 | 178 | 313 | 485 |

**Table S3. Overview of all simulations.**

| System | Total number of beads | Metadynamics | Metainference (SAXS) | Number of beads for SAXS calculation | Number of replicas | Total Simulation length [$\mu$s] ( simulation length per replica [ns] ) |
|---|---|---|---|---|---|---|
| *Di-ubiquitin* | | | | | | |
| **Martini 2.2** | 13569 | x | | | 112 | 48 (428) |
| **Increased P-W interaction** | 13569 | x | | | 112 | 69 (616) |
| **With SAXS** | 13569 | x | x | 355 | 112 | 65 (581) |
| **With SAXS(every time step)** | 13569 | *x* | x | 355 | 112 | 58 (517) |
| **Martini 3 beta** | 13569 | | | | 112 | 242 (2160) |
| *Tri-ubiquitin* | | | | | | |
| **Increased P-W interaction** | 12501 | x | | | 112 | 99 (884) |
| **With SAXS** | 12501 | x | x | 489 | 112 | 60 (536) |
| *Tetra-ubiquitin* | | | | | | |
| **Increased P-W interaction** | 15445 | x | | 652 | 112 | 60 (536) |
| **With SAXS** | 15445 | x | x | 652 | 112 | 63 (562) |
| *Nemo* | | | | | | |
| **With SAXS** | 33698 | *x* | x | 400 | 64 | 8 (125) |

**Table S4. Decomposition of the average interaction energy between ubiquitin pairs (residue 2-70).**

| | Coulomb interaction [kJ/mol] | Lennard Jones interaction [kJ/mol] | Total [kJ/mol] | Ratio Coulomb interaction [%] |
|---|---|---|---|---|
| *Di-ubiquitin* | | | | |
| **Ub(1-2)** | -4.3 ± 0.2 | -11.7 ± 0.9 | -16.0 ± 0.9 | 26.8 ± 2.0 |
| *Tri-ubiquitin* | | | | |
| **Ub(1-2)** | -6.8 ± 0.4 | -37.8 ± 2.2 | -44.6 ± 2.2 | 15.2 ± 1.1 |
| **Ub(2-3)** | -5.7 ± 0.3 | -36.1 ± 2.3 | -41.8 ± 2.4 | 13.7 ± 1.0 |
| **Ub(1-3)** | -0.8 ± 0.2 | -6.6 ± 1.7 | -7.5 ± 1.7 | 11.4 ± 3.7 |
| *Tetra-ubiquitin* | | | | |
| **Ub(1-2)** | -4.8 ± 0.3 | -33.9 ± 2.5 | -38.7 ± 2.5 | 12.5 ± 1.1 |
| **Ub(2-3)** | -4.6 ± 0.3 | -32.2 ± 2.2 | -36.7 ± 2.2 | 12.4 ± 1.1 |
| **Ub(3-4)** | -5.4 ± 0.3 | -54.7 ± 4.4 | -60.1 ± 4.4 | 9.0 ± 0.8 |
| **Ub(1-3)** | -0.2 ± 0.1 | -1.4 ± 0.6 | -1.7 ± 0.6 | 14.3 ± 7.6 |
| **Ub(2-4)** | -0.3 ± 0.1 | -1.6 ± 0.8 | -1.9 ± 0.8 | 13.9 ± 7.6 |
| **Ub(1-4)** | -0.2 ± 0.1 | -1.7 ± 1.0 | -1.9 ± 1.0 | 9.8 ± 7.5 |

**Table S5**. **Concentrations of substrates used for SAXS measurements.** Protein concentration (c) was determined by measuring the absorbance at 205 nm using specific absorbance for $NEMO_{258–350}$ C347S of 300990 $M^{-1}$ $cm^{-1}$. *The concentration of $Ub_4$ was corrected based on SEC-SLS.

| System | ratio | c (mg/ml) | corrected c* | NEMO / uM | $Ub_x$ / uM |
|---|---|---|---|---|---|
| **NEMO** | | 7.72 | | 702.5 | |
| **NEMO** | | 4.62 | | 420.4 | |
| **NEMO** | | 2.34 | | 212.9 | |
| **$Ub_3$** | | 11.17 | | | 435.2 |
| **$Ub_3$** | | 6.72 | | | 261.9 |
| **$Ub_3$** | | 3.41 | | | 133.0 |
| **$Ub_4$** | | 10.2 | 15.1 | | 441 |
| **$Ub_4$** | | 6.14 | 9.1 | | 265 |
| **$Ub_4$** | | 3.59 | 4.5 | | 131 |
| **NEMO:$Ub_3$** | 1.37:1 | 8.29 | | 279.4 | 203.5 |
| **NEMO:$Ub_3$** | 1.37:1 | 4.19 | | 141.0 | 102.8 |
| **NEMO:$Ub_3$** | 2.77:1 | 7.12 | | 351.2 | 126.9 |
| **NEMO:$Ub_3$** | 2.72:1 | 3.59 | | 175.6 | 64.5 |
| **NEMO:$Ub_4$** | 1.04:1 | 7.01 | 9.25 | 210.2 | 202.9 |
| **NEMO:$Ub_4$** | 1.04:1 | 3.54 | 4.67 | 106.4 | 102.2 |
| **NEMO:$Ub_4$** | 2.08:1 | 6.83 | 8.48 | 308.7 | 148.6 |
| **NEMO:$Ub_4$** | 2.06:1 | 3.04 | 4.27 | 154.3 | 75.1 |
| **NEMO:$Ub_4$** | 3.11:1 | 6.97 | 8.30 | 377.8 | 121.4 |
| **NEMO:$Ub_4$** | 3.11:1 | 3.48 | 4.15 | 188.9 | 60.7 |

**Table S6. Isothermal titration calorimetry (ITC) measurement of the interaction of NEMO with $Ub_3$ and $Ub_4$.** The numbers show the fitting of the data for the individual experiments in **a** and **b**, with N for stoichiometry, $K_D$ for dissociation constant, ΔH for enthalpy change and –TΔS for entropy change.

| System | N | $K_D$ (μM) | ΔH (kJ/mol) | -TΔS (kJ/mol) |
|---|---|---|---|---|
| *$Ub_3$ + Nemo* | | | | |
| run 1 | 1.99 ± 0.01 | 2.14 ± 0.26 | -16.7 ± 0.23 | -15.7 |
| run 2 | 1.85 ± 0.01 | 1.25 ± 0.14 | -18.0 ± 0.21 | -15.7 |
| run 3 | 1.68 ± 0.01 | 1.33 ± 0.11 | -18.9 ± 0.16 | -14.6 |
| **mean ± std error** | **1.84 ± 0.03** | **1.57 ± 0.28** | **-17.9 ± 0.64** | **-15.3 ± 0.4** |
| *$Ub_4$ + Nemo* | | | | |
| run 1 | 1.99 ± 0.01 | 4.66 ± 0.40 | -19.0 ± 0.23 | -11.4 |
| run 2 | 2.10 ± 0.01 | 4.12 ± 0.29 | -18.6 ± 0.17 | -12.1 |
| run 3 | 1.97 ± 0.02 | 3.61 ± 0.49 | -18.8 ± 0.32 | -12.2 |
| **mean ± std error** | **2.02 ± 0.04** | **4.13 ± 0.30** | **-18.8 ± 0.1** | **-11.9 ± 0.3** |

**Table S7. Determination of the molecular weight of NEMO, $Ub_3$, $Ub_4$, NEMO-$Ub_3$ and NEMO-$Ub_4$ using size exclusion chromatography (SEC) in combination with static light scattering (SLS).** Apart from indicated with "#" the conditions were 50 mM Tris.HCl pH 8, 300 mM NaCl. # 50 mM sodium phosphate pH 7, 50 mM NaCl (ITC conditions). * Peaks are not fully separated. Peak of complex "integrated".

| Protein | Peak 1 / ml (MWexp / kDa) | Peak 2 / mL (MWexp / kDa) | MWcalc /kDa | Remarks |
|---|---|---|---|---|
| **NEMO dimer** | 15.5 (**20.6**) | - | **22.0** | - |
| **$Ub_3$** | 16.6 (**24.1**) | - | **25.7** | - |
| **$Ub_4$** | 15.9 (**32.3**) | - | **34.2** | - |
| **NEMO:$Ub_3$ 1.4:1** | 14.6 (**39.9**) **(45.0)*** | 16.5 (**24.6**) | **47.7 (2:1)** | Excess $Ub_3$ |
| **NEMO:$Ub_3$ 2:1** | 14.7 (**41.1**) | 17.0 (**22.8**) | **47.7 (2:1)** | Free $Ub_3$ |
| **NEMO:$Ub_3$ 2:1#** | 14.2 (**42.2**) | - | **47.7 (2:1)** | - |
| **NEMO:$Ub_3$ 4:1** | 14.6 (**40.8**) | 15.3 (**24.1**) | **69.6 (4:1)** | Excess NEMO |
| **NEMO:$Ub_4$ 1.0:1** | 13.7 (**53.0**) | 15.8 (**32.7**) | **56.2 (2:1)** | Excess $Ub_4$ |
| **NEMO:$Ub_4$ 2.0:1** | 13.6 (**56.2**) | - | **56.2 (2:1)** | No extra peaks. |
| **NEMO:$Ub_4$ 4.1:1** | 13.6 (**53.2**) | 15.2 (**21.3**) | **78.2 (4:1)** | Excess NEMO |
| **NEMO:$Ub_4$ 2.5:1#** | 13.1 (**53.4**) | 14.6 (**16.5**) | **78.2 (4:1)** | Excess NEMO |

**Table S8. SPR steady state affinities of NEMO binding to immobilized ubiquitins.**

| Protein | $K_D$ (μM) |
|---|---|
| $Ub_3$ | 9.6 ± 3.3 |
| $Ub_4$ | 6.4 ± 1.4 |

$K_D$ values are reported as the mean ± SEM (n=2)
